# A flexible summary-based colocalization method with application to the mucin Cystic Fibrosis lung disease modifier locus

**DOI:** 10.1101/2021.08.06.455333

**Authors:** Fan Wang, Naim Panjwani, Cheng Wang, Lei Sun, Lisa J Strug

## Abstract

Mucus obstruction is a central feature in the Cystic Fibrosis (CF) airways. A genome-wide association study (GWAS) of lung disease by the CF Gene Modifier Consortium (CFGMC) identified a significant locus containing two mucin genes, *MUC20* and *MUC4*. Expression quantitative trait locus (eQTL) analysis using human nasal epithelial (HNE) from 94 CF Canadians in the CFGMC demonstrated *MUC4* eQTLs that mirrored the lung association pattern in the region, suggesting that *MUC4* expression may mediate CF lung disease. Complications arose, however, with colocalization testing using existing methods: the locus is complex and the associated SNPs span a 0.2Mb region with high linkage disequilibrium and evidence of eQTLs for multiple genes and tissues (heterogeneity). We previously developed the *Simple Sum* (*SS*), a powerful colocalization test in regions with heterogeneity, but SS assumed eQTLs to be present to achieve type I error control. Here we propose a two-stage SS (SS2) colocalization test that avoids *a prior* eQTL assumptions, accounts for multiple hypothesis testing and the composite null hypothesis and enables meta-analysis. We compare SS2 to published approaches through simulation and demonstrate type I error control for all settings with the greatest power in the presence of high LD and heterogeneity. Applying SS2 to the *MUC20/MUC4* CF lung disease locus with eQTLs from CF HNE revealed significant colocalization with *MUC4* (p = 1.71×10^−5^) rather than *MUC20*. The SS2 is a powerful method to inform the responsible gene(s) at a locus and guide future functional studies. SS2 has been implemented in the application LocusFocus (locusfocus.research.sickkids.ca).

## Introduction

Cystic Fibrosis (CF [MIM: 219700]) is a life-limiting genetic disease caused by mutations in the CF transmembrane conductance regulator (*CFTR* [MIM: 602421]). Multiple organs are affected in CF with variation in disease severity influenced by *CFTR* genotype, environmental factors and modifier genes.^1^ The majority of morbidity and mortality in CF results from lung disease which is heritable beyond the contributions of *CFTR*.^2^ Mucus pathology is a hallmark of CF airway disease, and thus the mucin family of genes have been hypothesized to contribute to lung disease severity in CF.^3^ A genome-wide association study (GWAS; *n* = 6365) of CF lung disease from the International CF Gene Modifier Consortium (ICFGMC) identified an associated locus on chromosome 3 in an intergenic region between two mucin genes, *MUC4* and *MUC20*, providing support for the mucin hypothesis but leading to uncertainty around the responsible gene(s) at the locus. Given the associated variants are not tagging protein coding variation, the assumption is that the associated locus is marking gene regulation.

Colocalization analysis using GWAS and gene expression quantitative trait locus (eQTL) summary statistics in a CF airway model can test this hypothesis and provide statistical support for the most probable gene(s) at the locus, however complications arise when we try to formally test colocalization at this locus using published tools.^4–15^ First, the null hypothesis of no colocalization is a composite null hypothesis that consists of several different null scenarios, e.g. the null scenario where there are significant GWAS and eQTL SNPs at a locus, but the two do not colocalize, or that there is a GWAS significant SNP but no eQTL at the locus (see Material and Methods for the four different null hypothesis scenarios). Type I error rate control under all possible null scenarios can be challenging for any method, especially in the presence of multiple hypothesis testing across multiple genes or tissues. Second, the associated SNPs at the *MUC4/MUC20* locus span a 0.2Mb region with evidence of allelic heterogeneity and high linkage disequilibrium (LD); these are factors that can substantially reduce the power of existing methods.^4,5^ Third, in our study, the participants from whom eQTLs are calculated overlap with a subset of the participants in the GWAS; this induces correlation between the eQTL and GWAS summary statistics that could lead to an increased false positive rate of colocalization depending on the degree of overlap.^16^ Lastly, our GWAS summary statistics are obtained from a meta-analysis with related individuals, further complicating LD adjustment in colocalization inference.

Several statistical methods have been developed to test for colocalization between GWAS and eQTL summary statistics, but none of the existing methods accommodate all four of these considerations. Bayesian colocalization approaches that are amenable to the use of summary statistics include eCAVIAR,^5^ COLOC,^4^ ENLOC,^9^ GWAS-PW,^6^ and COLOC2.^14^ These methods aim to identify whether there is a shared *causal* genetic variant that contributes to both the disease outcome and gene expression variation. Both eCAVIAR and ENLOC compute SNP-level posterior probabilities for being causal and a regional colocalization probability (RCP) by summing up the SNP-level probabilities. The RCP is computed for a locus that has been identified by GWAS, where the evidence under the composite null hypothesis is not explicitly calculated so it is unknown whether this method will have reliable operating characteristics more generally. COLOC, in contrast, computes the posterior probability incorporating subjective priors under five scenarios: four null scenarios and a specific alternative when one single causal variant is shared. Colocalization is concluded using a recommended value for the posterior colocalization probability (e.g. 0.8);^14^ explicit recommendations to control the false positive rate for multiple hypothesis testing is not available, to our knowledge. GWAS-PW extends COLOC by empirically estimating priors from the genome-wide data for the five scenarios to investigate genetic variants that influence a pair of traits and provides software functions to account for overlapping participants in the two studies.^6^ Similarly, COLOC2 incorporates features implemented in GWAS-PW and provides an updated version of COLOC.^14^ However, COLOC, GWAS-pw and COLOC2 all assume one single causal variant, and it has been shown that COLOC can have substantial loss in power when there are multiple, independent GWAS or eQTL signals at a locus (allelic heterogeneity).^8^ To gain additional power for a locus with multiple independent eQTL signals, Dobbyn et al.^14^ proposed a forward stepwise conditional analysis before conducting COLOC2 analyses. For each identified eQTL signal, COLOC2 integrates the GWAS result with eQTL evidence conditional on all the other eQTL signals. However, the conditional analysis requires individual-level data and can be computationally intensive.

There are several frequentist-based methods that also calculate colocalization evidence, for example the gene expression imputation approaches such as TWAS^13^ and prediXscan.^17^ These methods first use reference expression databases to define a set of genetic variants predictive of gene expression levels, then they impute gene expression levels in a study sample and test for association with a disease phenotype. The reliability of these methods depends on their prediction accuracy which can be limited.^18–20^ Extensions that enable the use of external summary statistics such as S-prediXscan,^7^ S-TWAS^13^ and S-multiXscan,^10^ and pre-computed parameters are available. However, in our case we already have gene expression data from a sample of CF patients in a relevant airway model. Integration methods based on Mendelian Randomization also *indirectly* address the question of colocalization. Two such approaches, SMR^11^ and Multi-SNP-based SMR (SMR-multi),^12^ derive their test statistics by assuming independence between GWAS and eQTL studies and use SNPs with eQTL p-values < 5 × 10^−8^ as instrumental variables to test whether gene expression differences cause the disease phenotype. SMR and SMR-multi restrict their analyses to regions with significant eQTLs, and they can accommodate meta-analysis but their robustness to related or overlapping samples is unknown. Lastly, JLIM^15^ evaluates whether there is a shared causal variant between eQTL and GWAS studies by developing a test statistic that contrasts the joint likelihood assuming one shared causal variant compared to that which assumes distinct causal variants between eQTL and GWAS. To calculate accurate colocalization p-values, JLIM requires individual-level gene expression data for permutation.

We previously developed an alternative frequentist and summary statistics-based colocalization method called Simple Sum (SS).^8^ It too does not address all of the four factors necessary for reliable colocalization analysis at our chr3q29 locus. SS performs well in the presence of linkage disequilibrium and allelic heterogeneity, however, at a given GWAS region with significant association but no eQTLs, the SS could have type I error inflation under this null scenario of no colocalization. Thus, we recommended an ad-hoc approach that restricted SS analyses to regions with significant eQTL evidence, similar to those imposed by other methods (e.g. SMR and SMR-multi). Here we extend the SS and propose a principled two-stage SS colocalization method (SS2) that controls type I error for the composite null hypothesis, even in the presence of multiple hypothesis testing across multiple genes or tissues, and remains powerful in the presence of LD and heterogeneity. The SS2 can also accommodate summary statistics calculated from related samples and meta-analysis, and we demonstrate the robustness of the method to the presence of a subset of overlapping samples between the two sets of summary statistics. The SS2 has been implemented in the application LocusFocus.^21^

For method comparison, we choose COLOC and COLOC2 from the class of Bayesian methods, and SMR and SMR-multi from the class of frequentist methods. Among the three Bayesian approaches that consider the issue of the composite null hypothesis (COLOC, GWAS-PW and COLOC2), COLOC has been shown to have better performance than GWAS-PW when a single causal variant is shared between the two studies,^15^ while GWAS-PW can have better performance when the two causal variants are distinct. We thus implement COLOC2 that incorporates features of both GWAS-PW and COLOC. Among the frequentist colocalization approaches that are applicable to summary statistics, S-prediXscan, S-TWAS and S-multiXscan use prediction weights inferred from a publicly available source (PredictDB Data Repository; Web Resources) to estimate the association between the predicted transcriptome and phenotypes, which can lead to biased or anti-conservative statistical inference.^18–20^ In our application, we already have expression from CF tissue (human nasal epithelial). Therefore, we implement SMR and SMR-multi which can be directly applied to summary statistics from any eQTL study without the need for training on a new sample to predict weights.

We first conduct extensive simulation studies to compare the proposed SS2 method with COLOC, COLOC2, SMR, and SMR-multi. We then apply these colocalization methods to the GWAS summary statistics from the *MUC4/MUC20* (chr3q29) CF lung function locus^22^ and the eQTL summary statistics, respectively, for *MUC20* and *MUC4* in CF primary human nasal epithelial (HNE). Finally, for completeness we extend the colocalization analyses to the genes within 1Mb of the locus using CF primary HNE, as well as other CF-related tissues from the genotype tissue expression project (GTEx).^23^ We demonstrate statistical support for *MUC4* as the responsible gene at the locus. The *MUC4* impact appears to be relevant in the lungs, both from GTEx and CF HNE, even after adjustment for multiple hypothesis testing of all 564 gene-by-tissue pairs evaluated at this locus.

## Material and Methods

### Notation and Model

For a locus of interest (e.g. chr3q29), here we assume that available information includes summary statistics from a phenotype-SNP GWAS association analysis and an expression-SNP eQTL analysis for a gene and tissue of interest (e.g. *MUC20* in human nasal epithelial), although this is generalizable to any SNP-level summary data. Assume there are *m* SNPs at the locus, let ***Z*** = (*Z*_1_, … *Z*_*j*_, …, *Z*_*m*_)′ be the vector containing the corresponding GWAS summary statistics and ***T*** = (*T*_1_, … *T*_*j*_, …, *T*_*m*_)′ be the vector containing the eQTL summary statistics, where *T*_*j*_ represents the eQTL evidence for SNP *j* in a specific gene-by-tissue pair, *j* = 1, …, *m*. ***T*** can be obtained from, for example, GTEx^23^ or one’s own expression study, although the latter requires additional care if the GWAS and eQTL study samples overlap, which we discuss below.

Our alternative of interest is that the GWAS signal and the eQTL signal coincide. As the complement to this alternative, the null hypothesis, *H*_0_, that there is no colocalization is composite, including four different scenarios:^4^

*H*_01_: no SNP-phenotype association and no eQTL,
*H*_02_: no SNP-phenotype association but eQTL present,
*H*_03_: SNP-phenotype association present but no eQTL,
*H*_04_: both SNP-phenotype association and eQTL present but occurring at two independent variants.

That is, *H*_0_ = *H*_01_ ∪ *H*_02_ ∪ *H*_03_ ∪ *H*_04_.

To test the composite *H*_0_, the Simple Sum test statistic is defined as

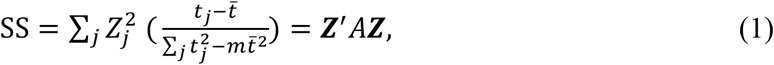

where 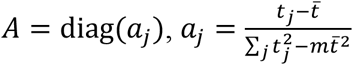, *j* = 1, …, *m*, and *t_j_* is a continuous eQTL evidence measure that can be defined as 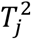 or −log_10_(eQTL p).^8^ In practice, 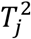 and −log_10_(eQTL p) produce SS p-values that provide the same colocalization conclusion. Under the null hypothesis scenarios, *H*_01_ ∪ *H*_02_, that there is no SNP-phenotype association, ***Z***~*N*(0, Σ) with the covariance matrix Σ capturing the LD structure of the region of interest (e.g. chr3q29).^24–28^ Thus, SS is distributed as 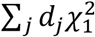 under *H*_01_ ∪ *H*_02_, where the *d*_*j*_’s are the eigenvalues of 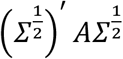. Implementing a one-sided test will then control the type I error not only under *H*_01_ or *H*_02_ but also under the null scenario of *H*_04_; the test statistic SS tends to be negative when the SNP-phenotype association and eQTL signals occur at two independent SNPs.^8^ However, this test could have type I error inflation under *H*_03_,^8^ and therefore we previously used caution to interpret colocalization findings when calculated from a small observed eQTL signal.

### SS2 Colocalization Test and Type I Error Control

Here we develop SS2 to test the composite null hypothesis that there is no colocalization, controlling the type I error rate under all four scenarios. SS2 is a two-stage testing procedure that analytically evaluates the eQTL evidence at the region of interest prior to conducting the colocalization analysis. We further extend the method for use with summary statistics obtained from meta-analysis and related individuals, and we investigate the robustness of SS2 to GWAS and eQTL summary statistics calculated using overlapping samples.

If we let the matrix *A* in equation (1) be the identity matrix, the SS test statistic is simplified to 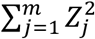, which has been used as a gene-based association test statistic.^29–31^ The distribution of this gene-based test statistic can be accessed by Monte Carlo simulation,^30^ by permutation,^29^ or it can also be derived analytically.^31^ The analytic distribution for this gene-based test was implemented in fastBAT^31^ where its power advantage was demonstrated over other gene-based tests that use the maximum of the 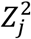’s, especially when there are multiple independent signals at a single locus. Here we replace the GWAS summary statistics used for a gene-based test, 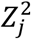’s, with the eQTL summary statistics 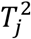’s, to evaluate the supporting evidence for an eQTL at the locus as *stage 1* of the SS2 test

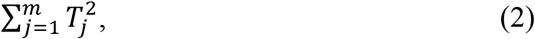

for a given gene in a tissue of interest.

Recall that under the null scenario of *H*_03_ where there is SNP-phenotype association but no eQTL, the original SS test has inflated type I error rate because the assumption of ***Z***~*N*(0, Σ) is violated. However, ***T***~*N*(0, Σ) in this case, so 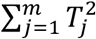 is distributed as 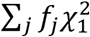, where *f*_*j*_’s are the eigenvalues of Σ.^31^ Thus, the SS2 stage 1 test rejects the null hypothesis of no eQTL if 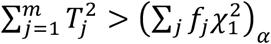, where 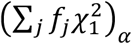 is the 1 − *α* quantile of 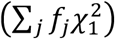. This stage 1 test alone controls the type I error of SS2 under *H*_03_ (and *H*_01_) and the type I error rates under the other null scenarios can be controlled by stage 2 of the SS2 colocalization test: If stage 1 of the SS2 test is significant, we then implement the SS test statistic of (1). That is, if ***Z***′ *A****Z*** > 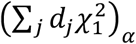, where 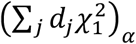 denotes the 1 − *α* quantile of 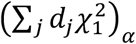, we conclude that there is colocalization of the eQTL and SNP-phenotype association.

In terms of the overall type I error rate control of the proposed SS2 two-stage test under the composite null, *H*_01_ ∪ *H*_02_ ∪ *H*_03_ ∪ *H*_04_, intuitively, under the null scenarios of *H*_01_ or *H*_03_ when there are no eQTLs, stage 1 already controls the false positives. Under the null scenarios of *H*_02_ or *H*_04_, even if the power of detecting the eQTL evidence is 100% in stage 1, stage 2 then provides the control of false positives. In the Supplemental Information, we show the independence between the stage 1 and stage 2 tests and demonstrate analytically the type I error rate control of SS2.

To ensure type I error rate control under the composite null hypothesis across all four different null scenarios, SS2 is conservative for certain null scenarios. For example, under *H*_01_ where there is no SNP-phenotype association or eQTL evidence, both stage 1 and stage 2 tests control the false positive rate at the nominal *α* level; by the independence of the two tests, the overall type I error rate is *α*^2^ (Table 2). This conservativeness is necessary so that the type I error rate under other null scenarios (e.g. *H*_03_) is not inflated. Furthermore, this conservativeness is necessary for conducting multiple colocalization hypothesis tests where the family of tests could consist of different null scenarios, which we discuss in the section on *Colocalization Testing in the Presence of Multiple Hypothesis Testing*.

**Table 1.**
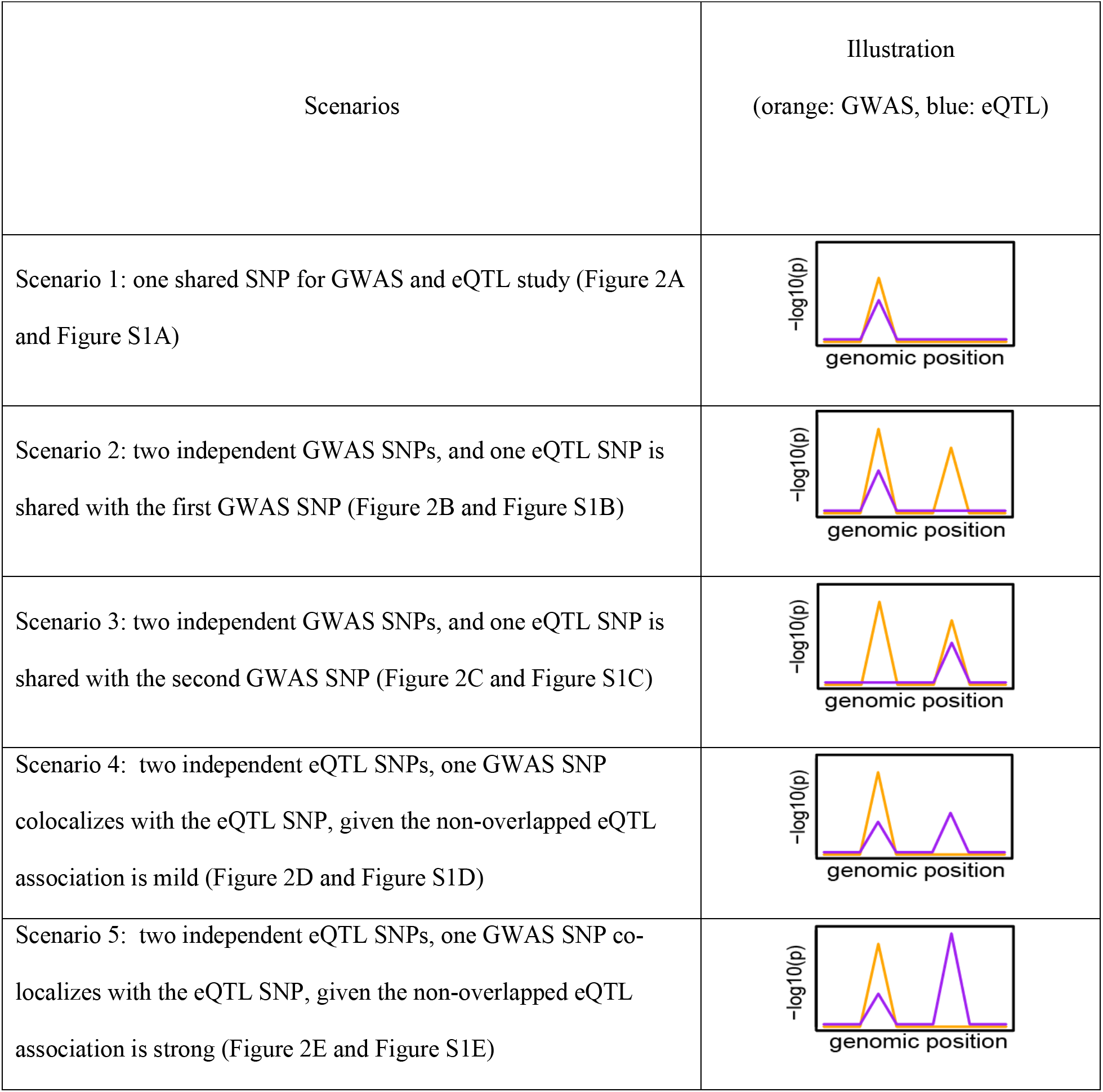
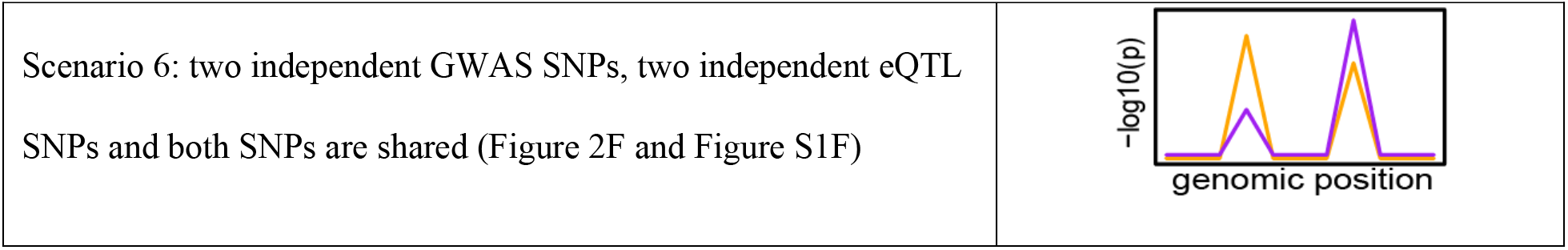
Overview and illustration of the six alternative scenarios we simulated to assess the power/true positive rate of different methods. The figures provide the general visualization of GWAS (orange line) and eQTL (blue line) colocalization patterns (on the −log10 p scale) in a region of interest.

**Table 2.**
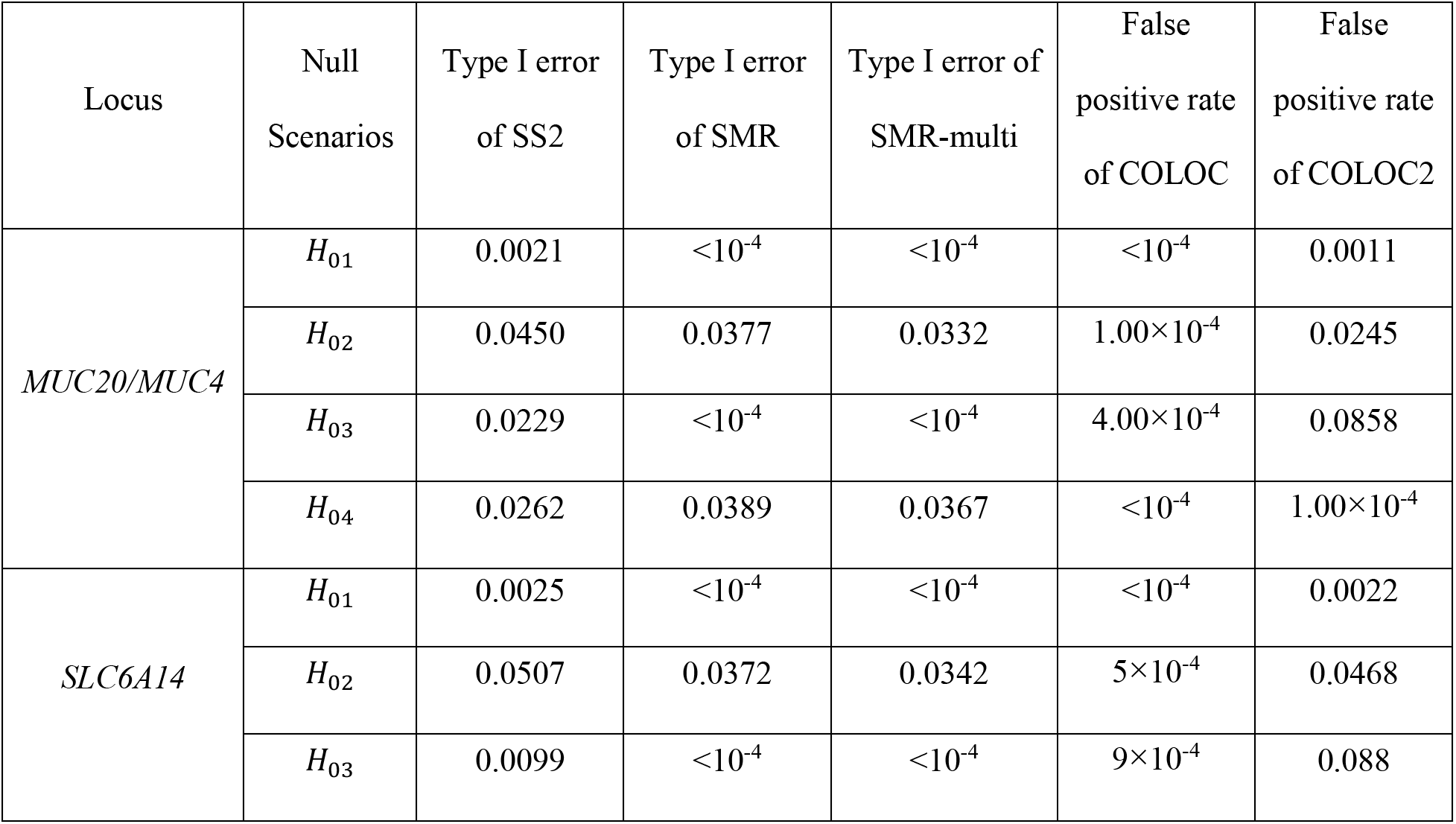
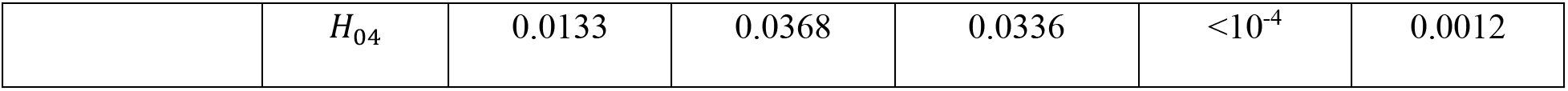
Empirical type I error rates of SS2, SMR and SMR-multi and false positive rates of COLOC2 and COLOC for a single hypothesis test. The LD pattern at the simulated region follows that at the *MUC20/MUC4* and *SLC6A14* locus, respectively. Each row corresponds to a specific null scenario when there is no co-localization. *H*_01_ represents the scenario when there are no SNP-phenotype associations and no eQTL; *H*_02_ represents the scenario when there are no SNP-phenotype associations but eQTLs are present; *H*_03_ represents the scenario where SNP-phenotype associations are present but no eQTL; *H*_04_ represents the scenario where both SNP-phenotype association and eQTLs are present, but occurring at two independent SNPs. For SS2, SMR^11^ and SMR-multi,^12^ the nominal type I error was set at *α* = 0.05. SMR and Multi-SNP-based SMR test (SMR-multi) are conducted under the default setting such that a SNP is picked if only if the eQTL p-value is less than 5×10^−8^. For COLOC2^14^ and COLOC,^4^ the false positive rates are calculated by applying the 0.8 threshold (as recommended by ^14^) for the colocalization posterior probability. In total, 10^4^ replications are simulated for each null scenario.

### Meta-analyses and Related Individuals

Many GWASs, such as our CF lung function GWAS, use a fixed or random effects meta-analysis to combine association evidence across multiple studies,^22,32^ and these multiple studies may contain related individuals. To implement a colocalization analysis using meta GWAS Z-scores, COLOC^4^ assumes unrelated individuals, while COLOC2^14^ incorporates features from GWAS-PW^6^ that consider sample overlap between studies but not sample relatedness within a study. SMR^11^ is derived based on the assumption that samples are independent between two studies, and its operating characteristics are unknown when there are related samples in a component study. Other methods such as eCAVIAR,^5^ ENLOC,^9^ SMR-multi,^12^ and JLIM^15^ compute their statistics under the assumption that GWAS Z-scores follow a multivariate normal distribution with covariance matrix Σ, where Σ is estimated assuming an independent sample using either one’s own data or external data such as that from the 1000 Genomes Project.^33^ However, accounting for the sample relatedness in the LD matrix, Σ, is important for valid colocalization analysis which we address below.

Assume the GWAS meta-analysis consists of *C* studies with sample sizes *n*_*c*_, *c* = 1, …, *C*. Let (*Z*_1,*j*_, …, *Z*_*C,j*_)′ and 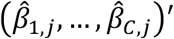 denote, respectively, the vector of Z scores and the vector of estimated effect sizes for SNP *j* from the *C* studies. Let *Z*_*meta,j*_ denote the *Z* score from the meta-analysis for SNP *j*, then

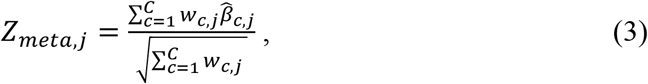

where *w*_*c,j*_ represents the weight for study *c*, which can take different forms.^32^ For the traditional fixed-effect approach, *w*_*c,j*_ is the inverse variance of 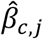 or estimated by 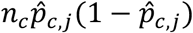, where 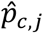 represents the estimated minor allele frequency for SNP *j* in study *c*. For the random-effect approach, the estimated between-study variance is incorporated into *w*_*c,j*_ to account for heterogeneity between studies.^32^

Based on equation (3), we can show that

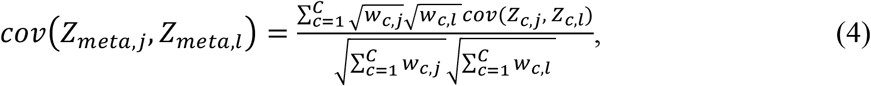

for SNPs *j* and *l*. When individuals in study *c* are independent of each other and one can assume a simple linear model for the SNP-phenotype association study, *cov*(*Z*_*c,j*_, *Z*_*c,l*_) = *r*_*c,jl*_, where *r*_*c,jl*_ is the standard Pearson correlation coefficient that represents the LD between SNPs *j* and *l* in study *c.*^5^ In this case, the covariance of meta-Z scores between the two SNPs is a weighted sum of study-specific LD measures. In the presence of related individuals, assume that the phenotype-SNP association test statistics are computed with a linear mixed effect model, where *Z* scores and their asymptotic covariance matrix can be written in a closed form that is equivalent to the estimation by generalized least squares (Supplemental Information). For the model that only contains the genotypes as predictors, 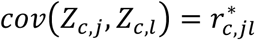, where 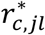 can be viewed as the pairwise Pearson correlation coefficient derived from the Cholesky-transformed genotype matrix. In the Supplemental Information, we provide a general form for *cov*(*Z*_*c,j*_, *Z*_*c,l*_) in the presence of additional covariates (e.g. age and sex) and discuss alternative ways to approximate *cov*(*Z*_*c,j*_, *Z*_*c,l*_) using for example the R package nlme or GMMAT.^34^

The eQTL summary statistics are typically obtained from a single study. If the eQTL summary statistics are also obtained from a meta-analysis, the covariance adjustment is also needed for conducting the stage 1 eQTL testing, following the principle in equation (4). This adjustment however does not influence the covariance adjustment in the stage 2 colocalization testing, where the inference is conditional on the observed eQTL evidence.

### Overlapping Samples between GWAS and eQTL Studies

The presence of overlapping samples can induce correlation between summary statistics even when there are no shared genetic effects, which may bias the model in favor of the alternative hypothesis.^6^ Let *Z*_1_ and *T*_1_ denote the summary statistics for a variant from a GWAS of sample size *n*_*GWAS*_ and an eQTL study for a given gene-by-tissue pair of sample size *n*_*eQTL*_, respectively,

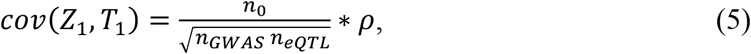

where *n*_0_ represents the number of overlapping samples; *ρ* represents the correlation between phenotypes for the overlapping samples due to non-genetic factors (e.g. shared environmental factors), and *ρ* equals 0 if there is no sample overlap.^6^ Equation (5) shows that the magnitude of the correlation between *Z*_1_ and *T*_1_, as expected, depends on the number of overlapping samples proportional to the sample size of each study, 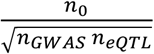, as well as the strength of the phenotypic correlation, *ρ*. Under the special case when two studies share the same sample (*n*_*GWAS*_ = *n*_*eQTL*_ = *n*_0_), the correlation between GWAS and eQTL summary statistics is completely determined by the phenotypic correlation, *cov*(*Z*_1_, *T*_1_) = *ρ*.^6^

Several published approaches have addressed the influence of overlapping samples in the context of two GWAS studies, and they have implemented decorrelation approaches that could be repurposed for our SS2 colocalization test.^6,16,35–37^ However, in our CF lung function chr3 application setting, the sample size of the lung GWAS study is 6,365 while the sample size of the eQTL study is 94. Thus, even under the extreme case that every individual included in the eQTL analysis overlaps with those in the GWAS study, 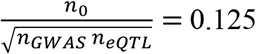, *cov*(*Z*_1_, *T*_1_) < 0.125, which would have a negligible impact on our inference. We demonstrate this empirically through a comprehensive simulation study (Table S1-S5).

### Colocalization Testing in the Presence of Multiple Hypothesis Testing

So far we have investigated the properties of SS2 when testing colocalization for a single gene in a specified tissue. In the present study, we are interested in determining whether the SNP-phenotype association evidence is colocalizing with gene expression of *MUC4* or *MUC20* in HNE, an established CF airway model.^3^ In fact, there are 50 genes annotated to a 1Mb region surrounding the top GWAS signal. Moreover, even though the *MUC4/MUC20* locus was identified as associated with CF lung disease, it would be of interest to investigate colocalization evidence in other tissues that may be affected in CF and for which the GTEx consortium provides eQTL summary statistics (GTEx V8).

To evaluate type I error rate control of the proposed SS2 colocalization test in the presence of multiple hypothesis testing, we consider the family-wise error rate (FWER). To maintain FWER control of SS2 under the composite null hypothesis for testing multiple genes and tissues, we implement stage 1 of the SS2 test of (2) for all the genes in each tissue and adjust the *α* for the total number of tests by Bonferroni correction, 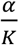. We then implement stage 2 of the SS2 test of (1) only for those *K*_2_ significant stage 1 eQTL tests and adjust for the corresponding multiple hypothesis testing by 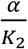.

The two-stage Bonferroni correction, 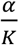 followed by 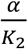, controls FWER at or below the nominal level. In the Supplemental Information, we show that when all *K* tests are under one of the four null scenarios, *H*_01_, *H*_02_, *H*_03_, or *H*_04_, the upper bound on the FWER of the two-stage SS2 test is *α*. In addition, when there is no SNP-phenotype association, the FWER of SS2 is also controlled at *α* even if the *K* tests consist of a mixture of *H*_01_ and *H*_02_. However, complications arise when there are SNP-phenotype associations at the locus but the *K* tests consist of a mixture of *H*_03_ and *H*_04_; this is of course a realistic scenario and has not been investigated before by us or by others.

To understand the challenge of FWER control in this mixture context, consider one simple scenario where the locus of interest was identified through GWAS as in the CF example, and there are two genes, *A* under *H*_03_ (no eQTL) and *B* under *H*_04_ (eQTL but not overlapping with GWAS). Let *α* denote the nominal significance level for the SS2 test and thus the stage 1 eQTL test is conducted at the 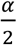 significance level for each gene. While testing the colocalization for gene *A* under *H*_03_, let *α*\* and 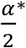 denote the empirical false positive rates of the stage 2 test at significance level *α* and 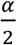 respectively. For gene *B*, let 1-*β* denote the power of the eQTL test in stage 1.

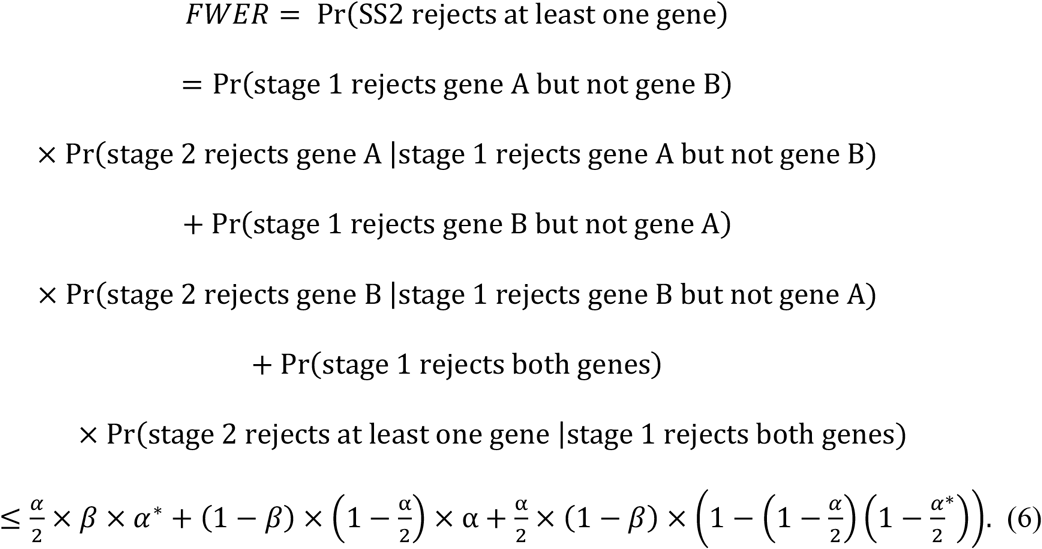

If 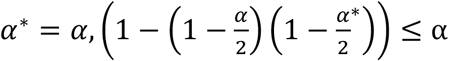, then equation (6)

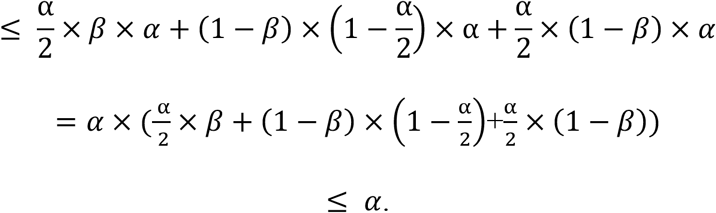

When the magnitude of *α*^∗^ is unknown, a crude upper bound for equation (6) can be specified by assuming the maximum value of *α*^∗^ to be 1, leading to equation (6) 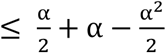; this is not necessarily less than or equal to *α* but provides a benchmark for the worst case scenario. In the Supplemental Information, we derive a crude upper bound that demonstrates the type I error is bounded but not necessarily at alpha. Our empirical studies below also show that the proposed two-stage SS2 testing procedure tends to be conservative, and we did not observe a single iteration with empirical FWER greater than the specified significance level, even under *H*_03_ (Table 3 and Table S8).

**Table 3.** Empirical family-wise error rates (FWERs) of SS2, SMR and SMR-multi and false positive rates of COLOC and COLOC2 for multiple genes. The height of the GWAS peak is set at 5.06 on the −log10p scale such that 10% power is achieved to detect the GWAS association at significance level of 10^−8^. In total, 600 genes are simulated based on the LD pattern at the *MUC20/MUC4* locus or the *SLC6A14* locus, respectively. Each row corresponds to a different proportion of genes that have eQTL association (0%, 20%, 40%, 60%, 80% and 100%). The eQTL peaks are randomly generated from 6 different intervals (50%-60%, 60%-70%, 70%-80%, 80%-90%, 90%-95%, 95%-100% power is achieved to detect the eQTL association at the significance level of 10^−8^) with probabilities according to the proportion of the −log10(maximum eQTL p-value) within each interval observed at the corresponding locus. None of the eQTL peak colocalizes with the GWAS peak for FWER evaluation. SMR and Multi-SNP-based SMR test (SMR-multi) are conducted under the default setting such that a SNP is picked if only if the eQTL p-value is less than 5×10^−8^. In total, 10^5^ replications are simulated to evaluate FWER of 0.05 and the false positive rates by applying the 0.8 threshold (as recommended by ^14^) for the colocalization posterior probability. The empirical FWER (or false positive rates for COLOC and COLOC2) is calculated by counting the proportions of 10^5^ replications where at least one gene has false colocalization claim.

| Locus | Proportion of genes with eQTL association but do not colocalize | FWER of SS2 | FWER of SMR | FWER of SMR-multi | False positive rate of COLOC | False positive rate of COLOC2 |
| --- | --- | --- | --- | --- | --- | --- |
| *MUC20/MUC4* | 0% | 0.0377 | 0.0010 | 0.0010 | 0.1307 | 0.0088 |
|  | 20% | 0.0294 | 0.0003 | 0.0002 | 0.1089 | 0.0023 |
|  | 40% | 0.0297 | 0.0002 | 0.0001 | 0.0854 | 0.0027 |
| | 60% | 0.0300 | $9.00 \times 10^{-5}$ | $5.00 \times 10^{-5}$ | 0.0606 | 0.0034 |
| | 80% | 0.0317 | $8.00 \times 10^{-5}$ | $4.00 \times 10^{-5}$ | 0.0351 | 0.0040 |
| | 100% | 0.0346 | $6.00 \times 10^{-5}$ | $5.00 \times 10^{-5}$ | 0.0043 | 0.0044 |
| *SLC6A14* | 0% | 0.0054 | 0.0008 | 0.0008 | 0.1626 | 0.0003 |
|  | 20% | 0.0016 | 0.0008 | 0.0004 | 0.1360 | 0.0049 |
|  | 40% | 0.0011 | 0.0004 | 0.0002 | 0.1103 | 0.0056 |
|  | 60% | 0.0010 | 0.0003 | 0.0002 | 0.0831 | 0.0070 |
|  | 80% | 0.0008 | 0.0002 | 0.0001 | 0.0542 | 0.0080 |
| | 100% | 0.0007 | 0.0001 | $9.00 \times 10^{-5}$ | 0.0234 | 0.0089 |

### Simulations

To evaluate the performance of the proposed SS2 colocalization test, we conduct extensive simulation studies using the LD pattern observed at the *MUC4*/*MUC20* locus and a second locus, for completeness, investigated previously, *SLC6A14*.^8^ For method comparison, we choose four alternative colocalization procedures, SMR, SMR-multi, COLOC, and COLOC2. We outline the simulation study design here and provide additional simulation details in the Supplemental Information.

#### Simulation for a Single Gene-by-Tissue Pair

We first consider a simulation study assessing colocalization at a locus between SNP-phenotype association p-values and SNP-expression (eQTL) p-values for a single gene in a given tissue type. Following the simulation procedure in ^5,8^, we focus on SNPs 0.1 Mb on either side of the top-associated SNP from the CF lung GWAS^22^ at the *MUC4*/*MUC20* locus and at the *SLC6A14* locus, respectively, to generate data for the two loci with different LD distributions. For type I error evaluation, we generate GWAS and eQTL summary statistics from a multivariate normal distribution based on the LD pattern of SNPs at each of the two loci and simulate null scenarios from *H*_01_ to *H*_04_ under the composite null hypothesis. Simulation details are provided in the Supplemental Information, and their corresponding parameter values are provided in Table S6.

We then assess the power of different methods by considering six alternative colocalization scenarios (Figure 2 and Figure S1), which go beyond the simple alternative when only one causal variant is shared by the GWAS and eQTL studies. Simulation details are provided in the Supplemental Information, and their corresponding parameter values are provided in Table S7.

#### Simulation for Multiple Gene-by-Tissue Pairs

We next evaluate the performance of the methods when studying colocalization evidence across many gene-by-tissue pairs. To be consistent with the presence of a GWAS signal and 564 gene-by-tissue pairs in the CF application, we simulate a locus, based on the LD pattern at the *MUC20/MUC4* locus, with one GWAS signal and 600 sets of independent eQTL summary statistics. We also considered 100, 200, 300, 400, or 500 colocalization tests simultaneously, varying the composition of the different alternative and null scenarios. Finally, we repeat the analysis for *SLC6A14*, a locus we studied previously ^8^ with a LD pattern different from *MUC20/MUC4*.

To evaluate the empirical FWER control, the simulated locus of interest has GWAS association evidence as in the CF application, while the eQTL summary statistics are simulated either under the null scenario of no eQTL (*H*_03_) or with an eQTL but does not colocalize with the GWAS signal (*H*_04_). When none (0%) of the 600 genes have eQTL evidence, all 600 tests are under *H*_03_. We also consider five different proportions of genes (20%, 40%, 60%, 80%, and 100%) with eQTL summary statistics but under *H*_04_. When these proportions are varied, the 600 tests are a mixture of *H*_03_ and *H*_04_, which is challenging in terms of type I error control as discussed earlier.

To evaluate power across different amounts of eQTL evidence while the locus has the same SNP-phenotype association evidence as in practice, we vary the colocalized eQTL evidence among the genes analyzed. In 5% of the 600 genes, we simulate the eQTL summary statistics that colocalize with the SNP-phenotype association evidence (i.e. simulated under the alternative). For the remaining 95% of genes, we simulate a proportion of the eQTL summary statistics to have no eQTL signal (under *H*_03_) while the remaining to have eQTL signals distinct from the SNP-phenotype association signal (under *H*_04_). We calculate power (or true positive rates for COLOC and COLOC2) by determining the proportion of 10^5^ simulated replications where at least one gene from the alternative is correctly identified.

#### Simulation for Overlapping Samples

In the presence of overlapping samples, the summary statistics for a variant from the GWAS and eQTL studies are correlated, which can lead to increased false positives in theory.^6^ To evaluate the practical impact of sample overlap on SS2, we consider the scenario where half or all of the participants whose genotypes used to compute the eQTL summary statistics are also included in the GWAS. Mimicking the scenario in the CF application, we simulate 100 participants in the eQTL study and to be conservative we include additional 1900 participants in the GWAS (i.e. the GWAS sample size is 2000 of which 100 overlap with the eQTL study). We evaluate the empirical type I error rate of SS2 under the composite null hypothesis from *H*_01_ to *H*_04_, with four different phenotypic correlations, 0.3, 0.5, 0.7, and 0.9. For comparison, we also demonstrate the type I error rate when there are no samples overlapping using the same simulation procedure. In addition, considering a fixed level of phenotypic correlation (0.5 or 0.9) we also demonstrate the empirical type I error rate control of SS2 as we vary sample size for the eQTL study, from 100, 200, 300, 400 to 500. Lastly, we simulate the scenario where all GWAS samples are overlapping with eQTL samples (both with 2000 individuals) and demonstrate the type I error rate of SS2 under the composite null hypothesis.

### EQTL Analysis

#### Sample Source and Collection

We conducted RNA-sequencing of human nasal epithelial (HNE) cells (*n* = 94) collected as part of the CF Canada Sick Kids Program in Individual CF Therapy (CFIT).^38^ The HNE samples are from CFIT participants collected using a 3-mm diameter sterile cytology brush (MP Corporation, Camarillo, CA) or Rhino-probe curette on either inferior turbinate. Sequencing was performed in two rounds with Illumina HiSeq 2000 and HiSeq 2500 platforms (Illumina Inc. San Diego, California, USA), respectively. The HiSeq 2000 round was sequenced with 25 million paired-end reads (49 base pairs in length) per sample, and the samples processed using the HiSeq 2500 platform had average library size of 35 mill paired-end reads (124 base pairs in length).

#### RNA-Seq Data Processing and Analysis

Quality of sequencing reads was assessed using FastQC (ver. 0.11.5; Web Resources) before and after trimming by Trim Galore (ver. 0.4.4; Web Resources). Processed reads were aligned to human reference genome hg38 with GENCODE comprehensive gene annotation (release 29) using STAR (ver. 2.5.4b).^39^ The reference genome included alternative haplotype contigs to account for the sequence diversity in the mucin gene locus. Expression quantification was performed by RNA-SeQC (ver. 2.0.0), which generated both read counts and normalized transcripts per million (TPM) measures.^40^ Normalized trimmed mean of M values (TMM) measures were obtained for a sub-sample of genes with ≥ 0.1 TPM and ≥ 6 read counts in more than 20% of the sample.^41^

Expression quantitative trait loci were analyzed by conducting differential gene expression analysis of the effect of SNP genotypes on TMM-normalized expression level. eQTL analysis was carried out using FastQTL (ver.2.0)^42^ with RNA-sequencing (RIN ≥ 7) of HNE from 94 Canadians with CF enrolled in the ICFGMC; these 94 individuals were also included in the CF lung GWAS. Additional covariates adjusted for in the model include the top 3 principal components, 15 probabilistic estimates of expression residuals (PEER) factors, study sites, sex, genotyping platform, RNA integrity number (RIN) and PTPRC/CD45 gene expression (immune cell content adjustment). R packages GENESIS (version 2.14.3) and peer (version 1.0) were used to generate genotype principal components and PEER factors, respectively.^43–45^

### Analysis of Colocalization at the *MUC4/MUC20* Locus

Genome-wide association analysis of CF lung function by the International Cystic Fibrosis Gene Modifier Consortium was carried out in 6,365 individuals (including 1,443 sib-pairs) with CF using a fixed-effect meta-analysis aggregating evidence from 13 subgroups.^22^ Among other loci, 24 genome-wide significant SNPs were identified and annotated between two mucin genes: *MUC4* and *MUC20* on chromosome 3.

To determine whether gene expression variation at the chr3q29 locus could influence lung disease severity and which genes the associated SNPs impact, we first focused on colocalization analysis for *MUC4* and *MUC20* given their biologic plausibility. For completeness, we subsequently expanded our analysis to investigate eQTLs of 50 genes in a 1Mb region on either side of the peak and in 14 CF-relevant tissues from GTEx V8. We focus on SNPs in a 0.1Mb region on either side of the lead GWAS SNP for colocalization analysis of all 564 gene-by-tissue pairs for which there is gene expression data.

To conduct the SS2 stage 1 eQTL test using (2), we computed the LD matrix for the eQTL analysis in HNE by calculating the Pearson correlation coefficient using the 94 CF independent samples, while for the eQTL studies in other CF-related tissues we use the GTEx resource. To apply the SS2 stage 2 colocalization test to the GWAS meta-summary statistics while accounting for sample relatedness, we estimated the LD matrix using equation (4), where the covariance of summary statistics for the sub-studies were calculated from Cholesky-transformed genotype data.

## Results

### Simulation Results

#### Simulation Results for a Single Gene-by-Tissue Pair

Table 2 demonstrates the empirical type I error rates of SS2, SMR^11^ and SMR-multi,^12^ and the false positive rates of COLOC^4^ and COLOC2^14^ for each of the four null scenarios of the composite null hypothesis, based on the LD pattern observed at the *MUC20/MUC4* locus or the *SLC6A14* locus. The SS2, SMR and SMR-multi all control the type I error rate at or below the nominal *α* = 0.05 significance level under all four null scenarios. When there are no eQTLs, under *H*_01_ or *H*_03_, SMR and SMR-multi are extremely conservative (the empirical *α* < 10^−4^). This is due to the recommended pre-screening step whereby colocalization analysis can only be conducted when there is an eQTL p-value less than 5×10^−8^ at the locus under investigation. Other eQTL p-value thresholds such as 0.01 or 0.05 could be adopted when conducting the SMR or SMR-multi, however, there is type I error inflation under *H*_03_ when one chooses to do so (Table S18).

Among all the methods, COLOC shows the most conservative results under the different null scenarios. The false positive rate for COLOC2 is controlled except for the *H*_03_ scenario. When there is SNP-phenotype association but no eQTLs, the empirical false positive rate is 0.09 for the nominal level of 0.05. The empirical false positive rate of COLOC2 decreases as one increases the sample size of the eQTL study (Table S17). For example, with 2000 participants in the GWAS, the empirical false positive rate of COLOC2 is below 0.05 when the sample size for the eQTL study is larger than 1000. However, in practice, the sample size of an eQTL study (e.g. GTEx)^23^ is usually much smaller than 1000, and in our CF study we only have 94 participants included in the eQTL analysis in HNE. We observed qualitatively similar results with the different LD patterns between the *MUC20* and *SLC6A14* loci.

The effect of either half or all of the participants in the eQTL study (100 individuals) overlapping with the GWAS study (2000 individuals) on the type I error rate of the SS2 is demonstrated in Tables S2-S5. The type I error rate of the SS2 remains controlled with increasing phenotypic correlation and increasing eQTL sample sizes, which demonstrates that the overlapping samples have minimal impact on the inference in practice.

We evaluate the power of SS2, SMR and SMR-multi at the *α* = 0.05 significance level and the true positive rate of COLOC2 and COLOC using a posterior probability of colocalization cut-off of 0.8 (Figure 2). Figure 2A illustrates the power under the simplest alternative Scenario 1, where one single SNP-phenotype association signal colocalizes with one single eQTL signal. In that case, COLOC2 has the highest true positive rate across different levels of eQTL evidence, but its false positive rate is inflated under *H*_03_ as demonstrated in Table 2. Among the methods that control the false positive rates across all null scenarios, namely SS2, SMR, SMR-multi, and COLOC, the proposed SS2 method is the most powerful.

When there are two GWAS peaks under the alternative Scenarios 2 and 3, (Figures 2B and 2C, respectively) and the eQTL evidence is weak (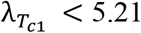 in Figure 2B and 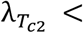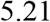 in Figure 2C; i.e. less than 30% power to detect the eQTL evidence at *α* = 0.05), SS2 has less power than COLOC2; this represents a trade-off to achieve type I error control under *H*_03_. However, SS2 is more powerful than the other three methods that do not have inflated false positive rates. SMR and SMR-multi are very conservative in this case because there are few SNPs with eQTL p-values less than 5×10^−8^. As the eQTL evidence gets stronger (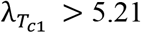 in Figure 2B and 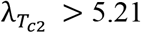 in Figure 2C; eQTL power greater than 30%), the power of SS2 could reach a similar level or even exceed the true positive rate of COLOC2. The power of SMR and SMR-multi are similar when there is only one eQTL SNP in the region (Figures 2A, 2B and 2C), and increase rapidly as the size of the eQTL signal increases. Under the alternative Scenario 3, when the eQTL evidence is strong (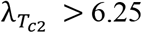; eQTL power greater than 70% power) and colocalizes with the second SNP-phenotype association peak (Figure 2C), SMR and SMR-multi are more powerful than SS2.

Figure 2D and 2E demonstrate the alternative Scenarios 4 and 5, respectively, when there are two independent eQTL SNPs but only one eQTL SNP colocalizes with the SNP-phenotype association. In this case, the SS2 is more powerful than all of the other methods across all levels of eQTL evidence considered. The power advantage of SS2 over other methods is especially notable when, between the two eQTL SNPs, the one with weaker signal colocalizes with the SNP-phenotype associated variant. Finally, under the alternative Scenario 6 when two independent GWAS SNPs colocalize with two independent eQTL SNPs, methods SS2, SMR, and SMR-multi show equally high power, while COLOC and COLOC2 have reduced power due to the allelic heterogeneity. Qualitatively similar results based on the LD pattern at the *SLC6A14* locus are evident in Figure S1.

#### Simulation Result for Multiple Genes-Tissue Pairs

Table 3 demonstrates the impact of multiple hypothesis testing on the family-wise error rate, where a locus with 600 genes and LD modeled after the *MUC20/MUC4* and *SLC6A14* loci were evaluated. Similar to the single hypothesis testing result in Table 2, SS2, SMR and SMR-multi control the FWER with the SMR-based tests being the most conservative. However, COLOC now has inflated false positive rates (> 0.08) for multiple scenarios, for example when 60% of genes at the locus have no eQTLs (*H*_03_) while the remaining 40% of genes have eQTLs but these are distinct from the SNP-phenotype association signal (*H*_04_). As the proportion of *H*_04_ genes decrease from 40% to 0% (or the proportion of *H*_03_ genes increase from 60% to 100%), the empirical false positive rate of COLOC increases from 0.08 to 0.13. Although COLOC2 shows an inflated false positive rate under *H*_03_ for our single hypothesis test investigation (Table 2), the false positive rate of COLOC2 is controlled after applying the algorithm implemented in GWAS-PW where the posterior probability is calculated based on the likelihood of all gene-by-tissue pairs.

We also investigated type I error rate control of the five methods when the number of genes tested at the locus was varied from 100 to 500 (Tables S8-S12). Overall, SS2, SMR and SMR-multi show conservative FWER (< 0.05; Table S8, S9 and S10 respectively). As the number of genes increases, the FWER of SS2 decreases when the LD is modeled after the *SLC6A14* locus and moderately increases when the LD is modeled after the *MUC20* locus. The increase in FWER tapers-off as the number of genes tested at the locus increase. This is because, with more genes passing the first stage test, the stage 2 colocalization test requires a more stringent significance level, resulting in the overall two-stage test being conservative enough to control the FWER. The false positive rate of COLOC increases as the number of genes evaluated increases, with inflation observed when the number of genes tested exceeds 200 (Table S11). This is due to the subjective choice of priors and cut-off (i.e. 0.8) for the colocalization posterior probability without explicit adjustment for multiple genes. In contrast, COLOC2 is conservative and stays conservative as one increases the total number of genes evaluated at a locus from 100 to 500 (Table S12).

Given the inflation in false positives for COLOC (Table S11), we compare power only between COLOC2, SMR, SMR-multi, and SS2. Keeping the eQTL evidence constant and with 600 genes, SS2 demonstrates the greatest power among the four methods; SMR has more power than SMR-multi (Table 4). The power of the four methods for testing 100 to 500 genes is provided in Tables S13-S16. SS2 shows consistently higher power compared to SMR when the number of genes is greater than 200, and SS2 shows higher power than SMR-multi, COLOC and COLOC2 across all gene numbers investigated. Interestingly, however, we observe different power trends with increasing numbers of genes analyzed at the locus. The power of SS2, SMR and SMR-multi decrease as the number of genes increases due to the Bonferroni correction, while the true positive rate of COLOC2 increases as the number of genes increases.

**Table 4.** Power of SS2, SMR and SMR-multi and true positive rate of COLOC2 for multiple genes. The LD pattern at the simulated region follows that at the *MUC20/MUC4* and *SLC6A14* locus, respectively. The height of the GWAS peak is set at 5.06 on the −log10p scale such that 10% power is achieved to detect the GWAS association at significance level of 10^−8^. Each row corresponds to a different range of the eQTL height for the 5% genes that have colocalization ([5.48, 5.73], [5.73, 5.98], [5.98, 6.25], [6.25, 6.57] and [6.57, 7.01]). The eQTL peaks are set with 5 different intervals such that 40%-50%, 50%-60%, 60%-70%, 70%-80%, 80%-90% power is achieved to detect the eQTL association at the significance level of 10^−8^. For the rest of 95% genes, there are eQTL evidence with mixture of null cases under *H*_03_ and *H*_04_ and details are demonstrated in the Supplemental Information. SMR and Multi-SNP-based SMR test (SMR-multi) are conducted under the default setting such that a SNP is picked if only if the eQTL p-value is less than 5×10^−8^. In total, 10^5^ replications are simulated to evaluate power at 0.05 significance level and the true positive rates by applying the 0.8 threshold (as recommended by ^14^) for the colocalization posterior probability. The power (or true positive rate for COLOC and COLOC2) is calculated by counting the proportions of 10^5^ replications where at least one gene is correctly identified with colocalization.

| Locus | EQTL height<br>for the 5%<br>genes have<br>colocalization | Power of<br>SS2 | Power of<br>SMR | Power of<br>SMR-multi | True positive rate of<br>COLOC2 |
| --- | --- | --- | --- | --- | --- |
| *MUC20/MUC4* | 5.48-5.73 | 0.8181 | 0.6826 | 0.6059 | 0.6218 |
|  | 5.73-5.98 | 0.8178 | 0.6935 | 0.6119 | 0.6355 |
|  | 5.98-6.25 | 0.8171 | 0.7033 | 0.6169 | 0.6447 |
|  | 6.25-6.57 | 0.8161 | 0.7144 | 0.6216 | 0.6507 |
|  | 6.57-7.01 | 0.8143 | 0.7270 | 0.6254 | 0.6536 |
| *SLC6A14* | 5.48-5.73 | 0.6855 | 0.6323 | 0.5531 | 0.6605 |
|  | 5.73-5.98 | 0.6992 | 0.6462 | 0.5614 | 0.6684 |
|  | 5.98-6.25 | 0.7076 | 0.6588 | 0.5679 | 0.6741 |
|  | 6.25-6.57 | 0.7138 | 0.6706 | 0.5745 | 0.6786 |
|  | 6.57-7.01 | 0.7184 | 0.6831 | 0.5817 | 0.6821 |

### Colocalization analysis at the *MUC4/MUC20* CF lung disease modifier locus

The eQTL p-values for *MUC4* visually colocalize with the GWAS p-values, which suggests that *MUC4* expression may mediate CF lung disease (Figure 1). We first assess the colocalization evidence for *MUC20* and *MUC4* using the CF lung GWAS meta-analysis summary statistics^22^ with eQTLs calculated from the HNE gene expression and genotype data of 94 individuals with CF (Table 5). For *MUC20* in the HNE, the stage 1 test does not provide evidence of an eQTL at the 5% level (uncorrected p-value = 0.083), suggesting that the eQTL evidence is not strong enough to move on to stage 2 colocalization analysis. For the *MUC4* eQTLs, both stage 1 and stage 2 tests are significant with p-value = 3.79×10^−7^ and 1.71×10^−5^, respectively, providing statistical evidence of colocalization consistent with the visualization (Figure 1).

**Figure 1.**
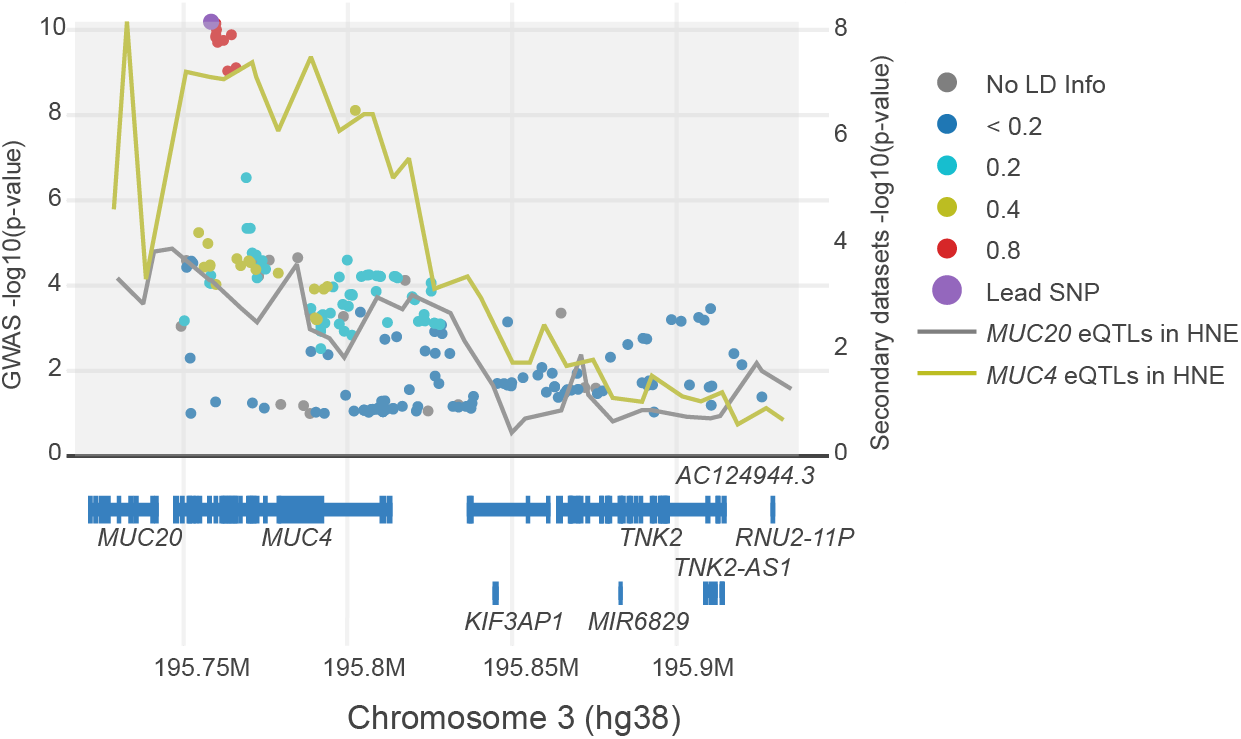
LocusFocus visualization of the CF lung disease GWAS summary statistics and expression quantitative trait locus (eQTL) summary statistics from primary human nasal epithelial (HNE) at the *MUC20/MUC4* locus. Overlay of p-values (on the −log_10_ scale) from the lung function GWAS (green/blue palette of colored dots) and eQTL association for *MUC20* and *MUC4* expression (colored lines) for HNE.

**Table 5.** Results of SS2, SMR, SMR-multi, COLOC, and COLOC2 applied to the *MUC20/MUC4* locus in primary human nasal epithelial (HNE) at chromosome 3. Colocalization analyses are conducted for all genes within a 1Mb region on either side of the peak lung GWAS associated variant and 14 CF-related tissues. In total, there are 564 gene-by-tissue pairs. Raw p-values and adjusted p-values by 564 gene-by-tissue pairs are both demonstrated for the SS2, SMR and SMR-multi. The eQTL evidence for conducting the SS2 is the eQTL p-value based on the −log10(eQTL p) scale for a specified gene and tissue. SMR and Multi-SNP-based SMR test (SMR-multi) are conducted under the default setting such that a SNP is picked if only if the eQTL p-value is less than 5×10^−8^. NAs are listed for *MUC20* since no SNP has eQTL p-value less than 5×10^−8^. For COLOC and COLOC2, the colocalization posterior probability (CLPP) is calculated, and a high posterior probability (> 0.8) suggests strong colocalization evidence. For COLOC2, we show both the CLPP calculated based on the likelihood from the single gene and tissue (CLPP1) and the CLPP calculated based on the likelihood from 564 gene-by-tissue pairs (CLPP2).

| Gene<br>and<br>tissue | SS2 stage 1 test |  | SS2 stage 2 test |  | SMR |  | SMR-multi |  | COLOC | COLOC2 |  |
| --- | --- | --- | --- | --- | --- | --- | --- | --- | --- | --- | --- |
|  | P-value | Adjusted<br>P-value | P-value | Adjusted<br>P-value | P-value | Adjusted<br>P-value | P-value | Adjusted<br>P-value | CLPP | CLPP1<sup>a</sup> | CLPP2 |
| *MUC20*<br>and<br>HNE | 0.083 | 1 | NA | NA<sup>b</sup> | NA | NA | NA | NA | 0.1057 | 0.0666 | 0.0005 |
| *MUC4*<br>and<br>HNE | $3.79 \times 10^{-7}$ | $2.10 \times 10^{-4}$ | $1.71 \times 10^{-5}$ | $1.95 \times 10^{-3}$ | $2.65 \times 10^{-3}$ | 0.379 | $2.07 \times 10^{-4}$ | 0.0296 | 0.7573 | 0.9482<sup>c</sup> | 0.0007 |

<sup>a</sup> This method has been shown with inflation in Table 3. CLPP1 > 0.8 suggests colocalization.
<sup>b</sup> The stage 1 SS2 test p-value (0.083) for *MUC20* and HNE does not pass the significant threshold (0.05) and therefore, the stage 2 SS2 test p-value is not applicable (NA).
<sup>c</sup> This method has been shown with inflation in Table 3.

To be comprehensive, we apply the SS2 to all genes annotated by GENCODE version 26 for hg38 GTEx V8 to the 1Mb region encompassing the peak CF lung-associated variant at the *MUC4/MUC20* GWAS locus. The colocalization evidence for each gene is calculated for the set of SNPs within 0.1Mb of the peak GWAS variant. Using the cross-tissue eQTLs from GTEx,^23^ we select the tissues that are relevant to CF and remove genes with low or no expression in a given tissue; this results in 564 gene-by-tissue pairs available for colocalization analysis. We apply the stage 1 set-based test on all the gene-by-tissue pairs, using a Bonferroni corrected significance level of 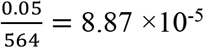. This results in 114 gene-by-tissue pairs providing evidence of significant eQTLs to move to the stage 2 test for colocalization. Stage 2 requires a significance level of 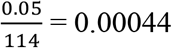 for each gene-by-tissue pair to conclude colocalization; 39 colocalization tests exceed this threshold. We present the SS2 cross-tissue and gene colocalization results in heatmaps (Figure 3A and Figure S2). For *MUC20* and *MUC4* we provide the stage 1 p-value adjusted by 564 gene-by-tissue pairs and the stage 2 p-value adjusted by 114 significant gene-by-tissue pairs in Table 5. *MUC4* remains significant after correction for the multiple tests at this locus, with stage 1 adjusted p-value of 2.10×10^−4^ and stage 2 adjusted p-value of 1.95×10^−3^, respectively.

For comparison, we also implement SMR, SMR-multi, COLOC, and COLOC2. Among the 564 gene-by-tissue pairs, SMR and SMR-multi are calculated on 143 gene-by-tissue pairs with top eQTL p-value less than 5×10^−8^, then applying a Bonferroni corrected significance level of 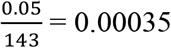. We provide both raw p-values and multiple testing adjusted p-values for SMR and SMR-multi analyses in Table 5. For *MUC20*, there are no SNPs with eQTL p-value smaller than 5×10^−8^ and therefore the SMR and SMR-multi test would not be applied. For *MUC4*, SMR provides a raw colocalization p-value of 2.65×10^−3^, but the multiple testing adjusted p-value = 0.379. SMR-multi demonstrates an association between gene-expression and the lung GWAS statistics with multiple testing adjusted colocalization p-value of 0.0296. Overall, the SMR and SMR-multi tests identified, respectively, 16 and 34 significant genes and tissues with colocalization evidence, which are shown in Figures 3B, 3C and Figure S2.

The colocalization posterior probability of COLOC2 for *MUC20* and *MUC4* are both small with 0.000472 and 0.000712, respectively. In this case, the empirical estimation of the prior for colocalization is low, which drags down the colocalization evidence for this locus. At this locus, there are no gene-by-tissue pairs with COLOC2 posterior probability higher than 0.8. In contrast, if we apply COLOC2 with the GWAS-PW algorithm ignoring the multiple hypothesis testing and focusing only on the likelihood from the single gene *MUC4*, the colocalization posterior probability is high (0.9482). However, the colocalization posterior probability for *MUC2*0 is low (0.0666; Table 5). A COLOC2 analysis based on the likelihood from each single gene-by-tissue pair at this locus provides 39 gene-by-tissue pairs with posterior probabilities higher than 0.8 (Figure S3).

The COLOC2 results at this locus are consistent with our simulation study, where COLOC2 has inflation of false positives when the GWAS-PW algorithm is implemented based on the likelihood from a single gene (Table 2). Yet, the test becomes over-conservative when priors are empirically estimated from the likelihood of multiple gene-by-tissue pairs (Table 3).

COLOC does not implement an approach to adjust for multiple hypothesis testing, but the empirical posterior probabilities of colocalization are < 0.8 (Table 5) for both *MUC20* (0.1057) and *MUC4* (0.7573). The results of COLOC applied to all 564 gene-by-tissue pairs are shown in Figure 3D and Figure S2.

## Discussion

The majority of associated genetic variants identified through GWAS fall in non-coding regions of the genome, thus the underlying mechanism by which the associated variants contribute to disease remains unclear but may point to gene regulation. The associations identified in the largest GWAS of CF lung disease to date^22^ are of no exception, with none of the five genome-wide significant loci tagging protein coding variation. One locus at chr3q29 is especially noteworthy as it encompasses *MUC4* and *MUC20,* members of a gene family that encode membrane-spanning ‘tethered’ mucins.^46–48^ These mucins prevent mucus penetration into the periciliary space and are present in the airway mucus, possibly contributing to mucociliary host defense. Mucus pathology is a defining characteristic of CF, with mucus hyperproduction and plugging, most notably in the CF airways.^3^ It has been presumed that mucus pathology is a downstream consequence of CFTR dysfunction,^8^ but GWAS identification at this locus suggests the possibility that polymorphisms impacting gene regulation of mucins may, themselves, impact the severity of CF lung disease. GWAS identifies loci. In contrast, colocalization analysis, integrating GWAS and gene expression data (represented by eQTLs), provides evidence that can connect the associated variants to gene expression changes, and pinpoints whether *MUC4*, *MUC20* or any of the other ~ 600 genes annotated to the locus could be putatively responsible.

Application of the SS2 to the CF lung disease associated locus at ch3q29 with eQTLs from CF HNE support that the associated lung disease variants colocalize with eQTLs for *MUC4*, prioritizing *MUC4* at the locus for further functional investigation. However, it should be noted that *MUC4* and *MUC20* are localized to a highly polymorphic region^22^ with several tandem repeats including a 48bp repeat region ranging from 7-19kb. The GWAS array data suggests a high frequency of large copy number variants around the clustered mucin region, but highly variable across individuals. This complex genomic context requires further consideration when differentiating between the two mucin genes at the locus.

There are several published colocalization methods, however the chr3q29 locus is in a region of high LD with evidence of allelic heterogeneity which poses challenges for existing procedures.^4–15^ Furthermore, the CF lung disease GWAS summary statistics are derived from a meta-analysis that includes sub-studies with related individuals; current methods cannot accommodate this scenario. We therefore developed the frequentist two-stage colocalization test, SS2. The SS2 integrates GWAS summary statistics with eQTL summary statistics across any number of gene-by-tissue pairs, is applicable when there are overlapping participants in the two studies and can be applied to GWAS summary statistics computed through meta-analysis, even with related individuals. Through simulation we demonstrate that the SS2 controls the type I error rate under the composite null hypotheses and is powerful in regions with high LD and allelic heterogeneity.

Bayesian colocalization approaches aim to identify a shared causal variant between two studies and to differentiate between distinct causal variants in LD. Similarly, Zhu et al implemented the Heidi test^11^ after the SMR test to further differentiate distinct causal variants in LD if the SMR test suggests significant association between two sets of summary statistics. Previous studies show that the statistical power of COLOC and Heidi for differentiating distinct causal variants decreases as the LD between causal variants increase.^11^ For a fair comparison, we calculated the FWER of SMR and SMR-multi tests without taking into account the results from the Heidi test since the SS2 test does not try to differentiate between distinct causal variants if they are in LD. In contrast, the SS2 aims to identify the association between two studies by leveraging the LD in the region and making inference based on the pattern similarity between summary statistics. Therefore, the SS2 can provide reliable inference even when the causal variant is not contained in the analysis set, as long as the LD pattern with the missing variant is retained.

The SS2 is a two-stage framework, designed to accomplish type 1 error control over the complex, composite null hypothesis. Although we implemented the 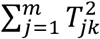 gene/set-based test as the first stage test of the SS2, there are many alternatives that could be implemented. These include versions with weighted sums of summary statistics, known as gene set analysis (GSA) tests or burden tests for rare variants.^49–51^ Summary statistics can also be decorrelated before being summed together, which is powerful under heterogeneity of effect sizes and variation between pairwise LD patterns.^49^ The stage 1 test implemented here has the same functional form as that used in VEGAS^30^ and fastBAT.^31^ This set-based test can be more powerful than a test which takes the maximum of the chi-squared test statistics in a region, an approach implemented in GATES^52^ and Pascal-Max,^53^ especially when there are multiple independent association signals.^31^ In contrast, SMR and SMR-multi use a p-value threshold (i.e. 5×10^−8^) to screen regions for analysis, presumably to ensure they are not using a weak instrument (this is similar to the approach implemented when the SS colocalization statistic was first defined).^8^ This stringent screening step results in power loss compared to SS2 when the eQTL association is only moderate which can be a function of several factors including sample size (Figure 2).

**Figure 2.**
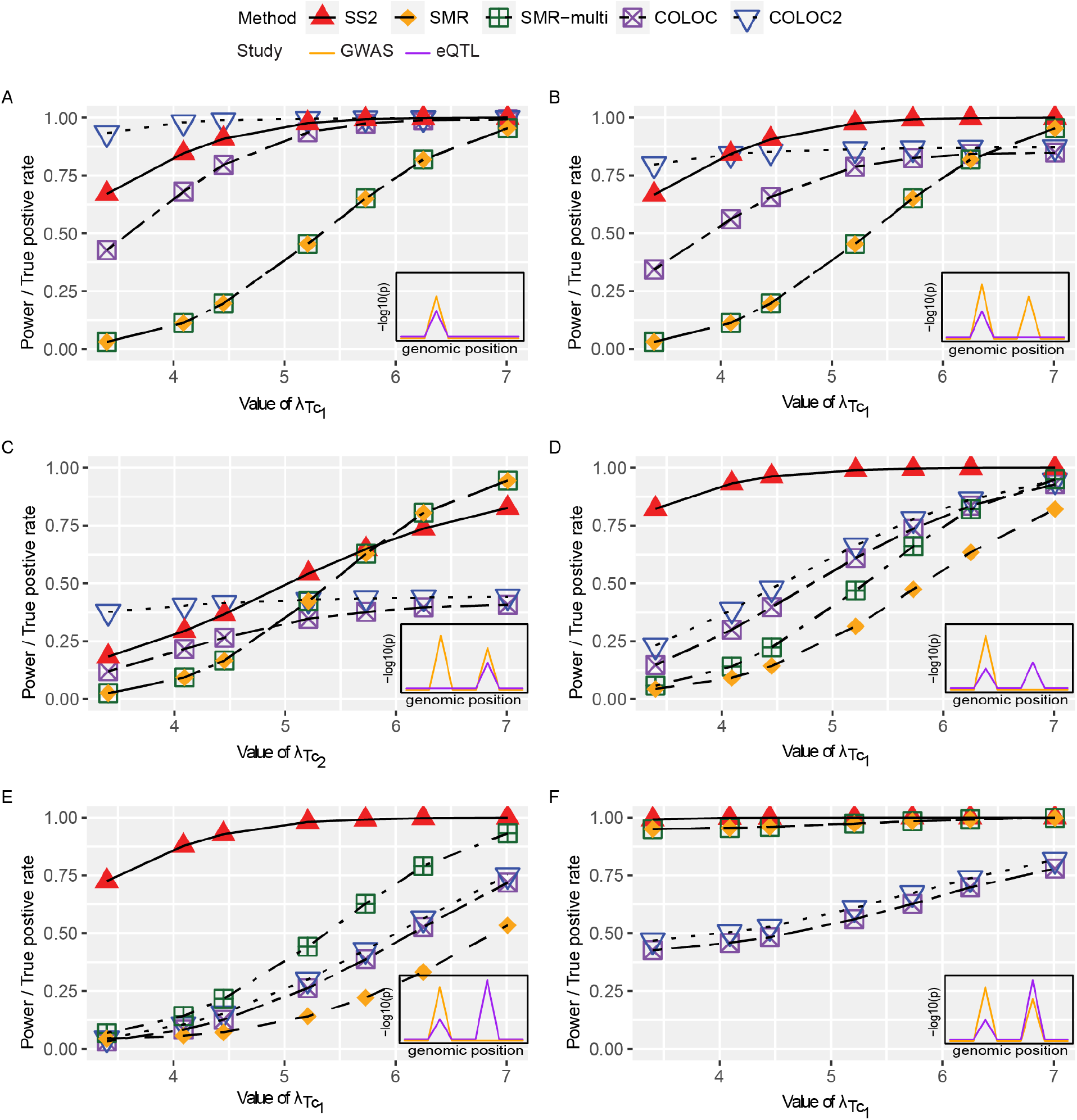
Empirical power of SS2, SMR and SMR-multi and true positive rates of COLOC and COLOC2 for testing a single gene for colocalization. The LD pattern at the simulated region follows that at the *MUC20/MUC4* locus at chromosome 3. For SS2, SMR^11^ and SMR-multi,^12^ the nominal type 1 error rate is set at *α* = 0.05. For COLOC2^14^ and COLOC,^4^ the false positive rates are controlled by applying the default 0.8 threshold (as recommended in ^14^) for the colocalization posterior probability. In total, 10^4^ replications are simulated to obtain the empirical power and true positive rates. The x-axis demonstrates the parameter values for 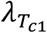 or 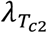 which is set to be 3.4, 4.09, 4.45, 5.21, 5.73 or 7.01 such that 0.01, 0.05, 0.1, 0.3, 0.5 or 0.9 power is achieved to detect the eQTL association at the significance level of 10^−8^. The inserted figure in the right bottom of each plot provides the general visualization of GWAS (orange line) and eQTL (purple line) colocalization patterns in a region of interest.

**Figure 3.**
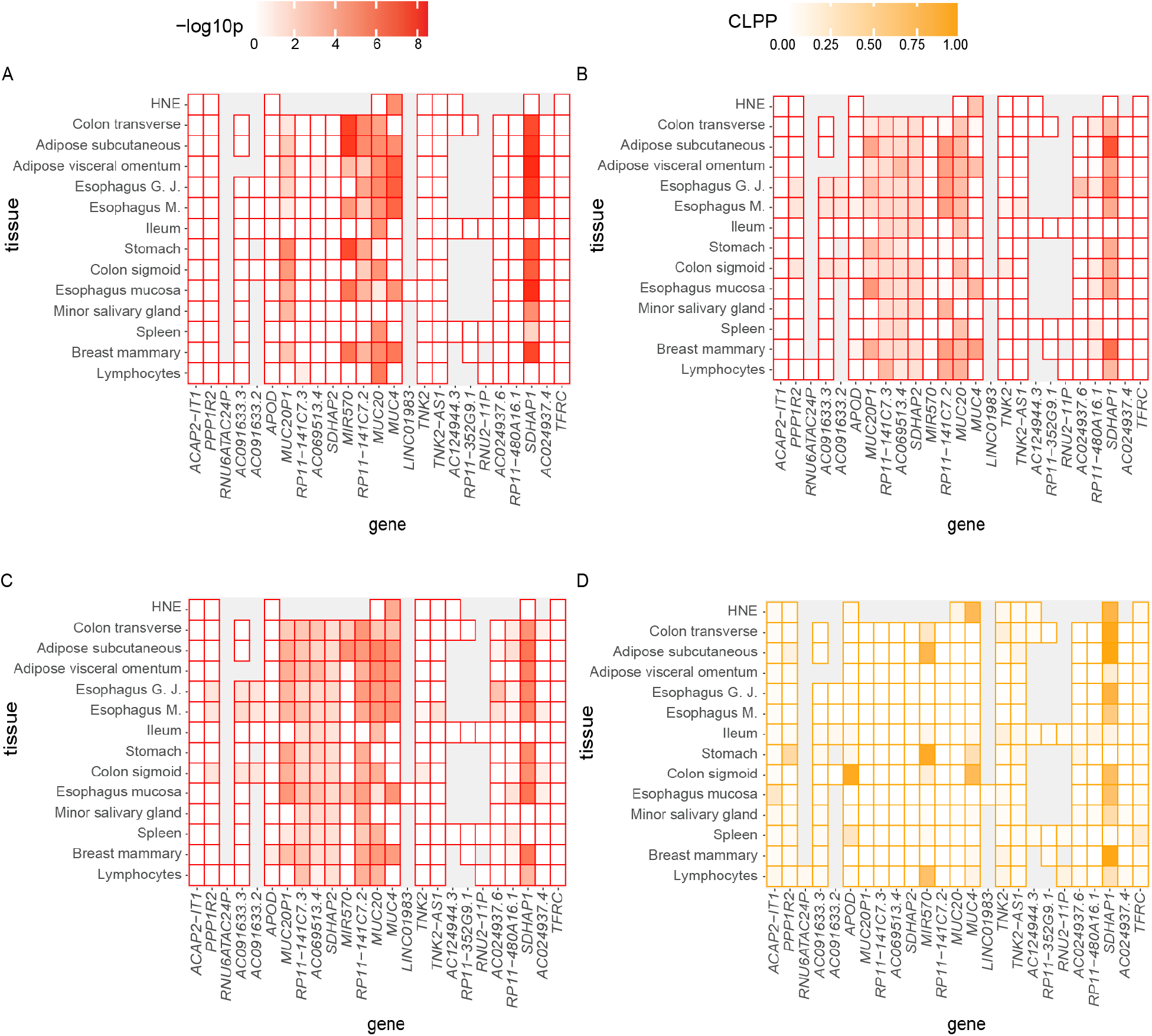
Heatmaps of colocalization evidence across genes and tissues at chromosome 3: (A) SS2, (B) SMR, (C) SMR-multi, and (D) COLOC. In each panel, each cell shows the colocalization evidence for the specified tissue and gene calculated from SNPs within 0.1Mb of the GWAS peak variant. Genes on the x-axis are ordered by their chromosomal positions. For illustration purposes, only the closest annotated genes centered around the GWAS peak variant are shown, and all genes analyzed within 1Mb around the GWAS peak variant is shown in Figure S2. Grey indicates insufficient expression levels attained for the gene in the tissue under study. (A): the color intensity corresponds to the SS2 colocalization evidence as measured by −log_10_(SS2 p-value), with red representing −log_10_(p) = 8.5 and white representing eQTL evidence for the corresponding gene and tissue does not pass the stage 1 test. (B) and (C): the color intensity corresponds to the SMR and SMR-multi colocalization evidence as measured by −log_10_(SMR p-value) and −log_10_(SMR-multi p-value), respectively; with red representing −log_10_(p) = 8.5 and white representing eQTL evidence for the corresponding gene and tissue does not pass the eQTL p-value threshold (5×10^−8^). (D): the color intensity corresponds to the COLOC colocalization evidence as measured by colocalization posterior probability (CLPP) ranging from 0 to 1. The eQTL analyses used for all gene/tissue pairs are those conducted by GTEx^23^ version 8 release, except for the HNE eQTLs. eQTL analyses in HNE is conducted using FastQTL^42^ with RNA-sequencing of HNE from 94 CF Canadians enrolled in the Canadian CF Gene Modifier study. Esophagus G.J represents Esophagus Gastroesophageal Junction; Esophagus M. represents Esophagus Muscularis; Ileum represents Small Intestine Terminal Ileum; Lymphocytes represents Cells EBV-transformed lymphocytes.

We modeled our simulation studies after the CF application and demonstrated that the SS2 has type I error rate control when 85 samples are included in both the GWAS and HNE eQTL studies. For other applications where there is a higher proportion of overlapping samples, the SS2 could have type I error inflation (Table S19) due to the correlation induced by the sample overlap. To address the effect of overlapping samples on statistical inference, several methods propose ways to estimate the correlation using summary statistics.^6,16,36,37^ In theory, one could then implement these approaches to decorrelate the summary statistics before applying the SS2 framework, although it was not necessary for our CF application as we showed.

When the eQTL summary statistics are replaced by GWAS evidence from a second phenotype, the SS2 framework enables the study of genetic overlap of the two traits. Similarly, the SS2 could assess colocalization using any SNP-level data including DNA methylation (meQTLs), protein QTLs (pQTLs) or metabolites (metQTLs). SS2 is implemented in a web-based colocalization tool, LocusFocus^21^ which enables integration of GWAS summary statistics with any secondary SNP-level dataset by using p-values and LD for the region of interest. The eQTL summary statistics from GTEx are made available for selection within the web server to test colocalization with tissues and genes from GTEx. All code and sample datasets are publicly available via GitHub under the MIT license.

## Supplemental Information

Supplemental information includes three figures (Figure S1-S3), 19 tables (Table S1-S19), and supplemental methods.

## Acknowledgments

We thank the patients, care providers and clinic coordinators at CF Centers throughout Canada for their contributions to the CF Patient Registry and Canadian CF Gene Modifier Study. We would like to thank the CF Canada-Sickkids Program for Individualized Therapy (CFIT) for generating the gene expression data. Funding was provided by Cystic Fibrosis Foundation STRUG17PO; Natural Sciences and Engineering Research Council of Canada (RGPIN-2015-03742, RGPIN-04934, RGPAS-522594), and the Canadian Institutes of Health Research (FRN-167282, FRN-310732), Cystic Fibrosis Canada (2626); and by the Government of Canada through Genome Canada (OGI-148) and supported by a grant from the Government of Ontario. The funders of the study play no role in study design, data collection and analysis, decision to publish or preparation of the manuscript. FW is a trainee of the CANSSI-Ontario STAGE training program at the University of Toronto.

## Web Resources

LocusFocus, https://locusfocus.research.sickkids.ca

GTEx Portal (release v.8), http://www.gtexportal.org

FastQC (ver. 0.11.5), https://www.bioinformatics.babraham.ac.uk/projects/fastqc/

Trim Galore (ver. 0.4.4), https://www.bioinformatics.babraham.ac.uk/projects/trim_galore/

PredictDB Data Repository, http://predictdb.org

## Data and Code Availability

Summary statistics from the GWAS are available at http://lab.research.sickkids.ca/strug/publications-software/.

R scripts to enable the extensions here can be found at https://github.com/FanWang0216/SimpleSum2Colocalization and https://github.com/naim-panjwani/LocusFocus.

Access to the RNA-sequencing data from the nasal epithelial are available through the CF Canada-SickKids Program for Individualized Therapy Biobank http://lab.research.sickkids.ca/cfit/.

## References

1. Cutting, G.R. (2010). Modifier genes in Mendelian disorders: the example of cystic fibrosis. Annals of the New York Academy of Sciences 1214, 57.

2. Vanscoy, L.L., Blackman, S.M., Collaco, J.M., Bowers, A., Lai, T., Naughton, K., Algire, M., McWilliams, R., Beck, S., and Hoover-Fong, J. (2007). Heritability of lung disease severity in cystic fibrosis. American journal of respiratory and critical care medicine 175, 1036–1043.

3. Kreda, S.M., Davis, C.W., and Rose, M.C. (2012). CFTR, mucins, and mucus obstruction in cystic fibrosis. Cold Spring Harb Perspect Med 2, a009589.

4. Giambartolomei, C., Vukcevic, D., Schadt, E.E., Franke, L., Hingorani, A.D., Wallace, C., and Plagnol, V. (2014). Bayesian Test for Colocalisation between Pairs of Genetic Association Studies Using Summary Statistics. PLOS Genetics 10, e1004383.

5. Hormozdiari, F., van de Bunt, M., Segrè, A.V., Li, X., Joo, J.W.J., Bilow, M., Sul, J.H., Sankararaman, S., Pasaniuc, B., and Eskin, E. (2016). Colocalization of GWAS and eQTL Signals Detects Target Genes. Am J Hum Genet 99, 1245–1260.

6. Pickrell, J.K., Berisa, T., Liu, J.Z., Ségurel, L., Tung, J.Y., and Hinds, D.A. (2016). Detection and interpretation of shared genetic influences on 42 human traits. Nature Genetics 48, 709–717.

7. Barbeira, A.N., Dickinson, S.P., Bonazzola, R., Zheng, J., Wheeler, H.E., Torres, J.M., Torstenson, E.S., Shah, K.P., Garcia, T., Edwards, T.L., et al. (2018). Exploring the phenotypic consequences of tissue specific gene expression variation inferred from GWAS summary statistics. Nature Communications 9, 1825.

8. Gong, J., Wang, F., Xiao, B., Panjwani, N., Lin, F., Keenan, K., Avolio, J., Esmaeili, M., Zhang, L., He, G., et al. (2019). Genetic association and transcriptome integration identify contributing genes and tissues at cystic fibrosis modifier loci. PLOS Genetics 15, e1008007.

9. Wen, X., Pique-Regi, R., and Luca, F. (2017). Integrating molecular QTL data into genome-wide genetic association analysis: Probabilistic assessment of enrichment and colocalization. PLOS Genetics 13, e1006646.

10. Barbeira, A.N., Pividori, M., Zheng, J., Wheeler, H.E., Nicolae, D.L., and Im, H.K. (2019). Integrating predicted transcriptome from multiple tissues improves association detection. PLOS Genetics 15, e1007889.

11. Zhu, Z., Zhang, F., Hu, H., Bakshi, A., Robinson, M.R., Powell, J.E., Montgomery, G.W., Goddard, M.E., Wray, N.R., Visscher, P.M., et al. (2016). Integration of summary data from GWAS and eQTL studies predicts complex trait gene targets. Nature Genetics 48, 481–487.

12. Wu, Y., Zeng, J., Zhang, F., Zhu, Z., Qi, T., Zheng, Z., Lloyd-Jones, L.R., Marioni, R.E., Martin, N.G., Montgomery, G.W., et al. (2018). Integrative analysis of omics summary data reveals putative mechanisms underlying complex traits. Nature Communications 9, 918.

13. Gusev, A., Ko, A., Shi, H., Bhatia, G., Chung, W., Penninx, B.W.J.H., Jansen, R., de Geus, E.J.C., Boomsma, D.I., Wright, F.A., et al. (2016). Integrative approaches for large-scale transcriptome-wide association studies. Nature Genetics 48, 245–252.

14. Dobbyn, A., Huckins, L.M., Boocock, J., Sloofman, L.G., Glicksberg, B.S., Giambartolomei, C., Hoffman, G.E., Perumal, T.M., Girdhar, K., Jiang, Y., et al. (2018). Landscape of Conditional eQTL in Dorsolateral Prefrontal Cortex and Co-localization with Schizophrenia GWAS. Am J Hum Genet 102, 1169–1184.

15. Chun, S., Casparino, A., Patsopoulos, N.A., Croteau-Chonka, D.C., Raby, B.A., De Jager, P.L., Sunyaev, S.R., and Cotsapas, C. (2017). Limited statistical evidence for shared genetic effects of eQTLs and autoimmune-disease-associated loci in three major immune-cell types. Nat Genet 49, 600–605.

16. LeBlanc, M., Zuber, V., Thompson, W.K., Andreassen, O.A., Frigessi, A., Andreassen, B.K., Schizophrenia, and Bipolar Disorder Working Groups of the Psychiatric Genomics, C. (2018). A correction for sample overlap in genome-wide association studies in a polygenic pleiotropy-informed framework. BMC Genomics 19, 494.

17. Gamazon, E.R., Wheeler, H.E., Shah, K.P., Mozaffari, S.V., Aquino-Michaels, K., Carroll, R.J., Eyler, A.E., Denny, J.C., Nicolae, D.L., Cox, N.J., et al. (2015). A gene-based association method for mapping traits using reference transcriptome data. Nature Genetics 47, 1091–1098.

18. Fryett, J.J., Morris, A.P., and Cordell, H.J. (2020). Investigation of prediction accuracy and the impact of sample size, ancestry, and tissue in transcriptome-wide association studies. Genetic Epidemiology 44, 425–441.

19. Wainberg, M., Sinnott-Armstrong, N., Mancuso, N., Barbeira, A.N., Knowles, D.A., Golan, D., Ermel, R., Ruusalepp, A., Quertermous, T., Hao, K., et al. (2019). Opportunities and challenges for transcriptome-wide association studies. Nat Genet 51, 592–599.

20. Wang, S., McCormick, T.H., and Leek, J.T. (2020). Post-prediction inference. BioRxiv.

21. Panjwani, N., Wang, F., Wang, C., He, G., Mastromatteo, S., Bao, A., Gong, J., Rommens, J.M., Sun, L., and Strug, L.J. (2020). LocusFocus: A web-based colocalization tool for the annotation and functional follow-up of GWAS. bioRxiv, 2020.2001.2002.891291.

22. Corvol, H., Blackman, S.M., Boëlle, P.-Y., Gallins, P.J., Pace, R.G., Stonebraker, J.R., Accurso, F.J., Clement, A., Collaco, J.M., Dang, H., et al. (2015). Genome-wide association meta-analysis identifies five modifier loci of lung disease severity in cystic fibrosis. Nature Communications 6, 8382.

23. Lonsdale, J., Thomas, J., Salvatore, M., Phillips, R., Lo, E., Shad, S., Hasz, R., Walters, G., Garcia, F., Young, N., et al. (2013). The Genotype-Tissue Expression (GTEx) project. Nature Genetics 45, 580–585.

24. He, X., Fuller, C.K., Song, Y., Meng, Q., Zhang, B., Yang, X., and Li, H. (2013). Sherlock: detecting gene-disease associations by matching patterns of expression QTL and GWAS. The American Journal of Human Genetics 92, 667–680.

25. Nica, A.C., Montgomery, S.B., Dimas, A.S., Stranger, B.E., Beazley, C., Barroso, I., and Dermitzakis, E.T. (2010). Candidate causal regulatory effects by integration of expression QTLs with complex trait genetic associations. PLoS Genet 6, e1000895.

26. Yanai, I., Benjamin, H., Shmoish, M., Chalifa-Caspi, V., Shklar, M., Ophir, R., Bar-Even, A., Horn-Saban, S., Safran, M., and Domany, E. (2005). Genome-wide midrange transcription profiles reveal expression level relationships in human tissue specification. Bioinformatics 21, 650–659.

27. Kryuchkova-Mostacci, N., and Robinson-Rechavi, M. (2017). A benchmark of gene expression tissue-specificity metrics. Briefings in bioinformatics 18, 205–214.

28. Howie, B., Fuchsberger, C., Stephens, M., Marchini, J., and Abecasis, G.R. (2012). Fast and accurate genotype imputation in genome-wide association studies through pre-phasing. Nature Genetics 44, 955–959.

29. Sun, L., Rommens, J.M., Corvol, H., Li, W., Li, X., Chiang, T.A., Lin, F., Dorfman, R., Busson, P.F., Parekh, R.V., et al. (2012). Multiple apical plasma membrane constituents are associated with susceptibility to meconium ileus in individuals with cystic fibrosis. Nat Genet 44, 562–569.

30. Liu, J.Z., McRae, A.F., Nyholt, D.R., Medland, S.E., Wray, N.R., Brown, K.M., Hayward, N.K., Montgomery, G.W., Visscher, P.M., Martin, N.G., et al. (2010). A Versatile Gene-Based Test for Genome-wide Association Studies. The American Journal of Human Genetics 87, 139–145.

31. Bakshi, A., Zhu, Z., Vinkhuyzen, A.A.E., Hill, W.D., McRae, A.F., Visscher, P.M., and Yang, J. (2016). Fast set-based association analysis using summary data from GWAS identifies novel gene loci for human complex traits. Scientific Reports 6, 32894.

32. Han, B., and Eskin, E. (2011). Random-effects model aimed at discovering associations in meta-analysis of genome-wide association studies. Am J Hum Genet 88, 586–598.

33. Auton, A., Abecasis, G.R., Altshuler, D.M., Durbin, R.M., Abecasis, G.R., Bentley, D.R., Chakravarti, A., Clark, A.G., Donnelly, P., Eichler, E.E., et al. (2015). A global reference for human genetic variation. Nature 526, 68–74.

34. Chen, H., Conomos, M.P., and Chen, M.H. (2019). Package ‘GMMAT’.

35. Lin, D.-Y., and Sullivan, P.F. (2009). Meta-Analysis of Genome-wide Association Studies with Overlapping Subjects. The American Journal of Human Genetics 85, 862–872.

36. Province, M.A., and Borecki, I.B. (2013). A correlated meta-analysis strategy for data mining “OMIC” scans. Pac Symp Biocomput, 236–246.

37. Zhu, X., Feng, T., Tayo, Bamidele O., Liang, J., Young, J.H., Franceschini, N., Smith, Jennifer A., Yanek, Lisa R., Sun, Yan V., Edwards, Todd L., et al. (2015). Meta-analysis of Correlated Traits via Summary Statistics from GWASs with an Application in Hypertension. The American Journal of Human Genetics 96, 21–36.

38. Eckford, P.D., McCormack, J., Munsie, L., He, G., Stanojevic, S., Pereira, S.L., Ho, K., Avolio, J., Bartlett, C., and Yang, J.Y. (2019). The CF Canada-Sick Kids Program in individual CF therapy: A resource for the advancement of personalized medicine in CF. Journal of Cystic Fibrosis 18, 35–43.

39. Dobin, A., Davis, C.A., Schlesinger, F., Drenkow, J., Zaleski, C., Jha, S., Batut, P., Chaisson, M., and Gingeras, T.R. (2013). STAR: ultrafast universal RNA-seq aligner. Bioinformatics 29, 15–21.

40. DeLuca, D.S., Levin, J.Z., Sivachenko, A., Fennell, T., Nazaire, M.-D., Williams, C., Reich, M., Winckler, W., and Getz, G. (2012). RNA-SeQC: RNA-seq metrics for quality control and process optimization. Bioinformatics 28, 1530–1532.

41. Robinson, M.D., and Oshlack, A. (2010). A scaling normalization method for differential expression analysis of RNA-seq data. Genome biology 11, 1–9.

42. Ongen, H., Buil, A., Brown, A.A., Dermitzakis, E.T., and Delaneau, O. (2016). Fast and efficient QTL mapper for thousands of molecular phenotypes. Bioinformatics 32, 1479–1485.

43. Gogarten, S.M., Sofer, T., Chen, H., Yu, C., Brody, J.A., Thornton, T.A., Rice, K.M., and Conomos, M.P. (2019). Genetic association testing using the GENESIS R/Bioconductor package. Bioinformatics 35, 5346–5348.

44. Conomos, M.P., Miller, M.B., and Thornton, T.A. (2015). Robust inference of population structure for ancestry prediction and correction of stratification in the presence of relatedness. Genetic epidemiology 39, 276–293.

45. Stegle, O., Parts, L., Piipari, M., Winn, J., and Durbin, R. (2012). Using probabilistic estimation of expression residuals (PEER) to obtain increased power and interpretability of gene expression analyses. Nature protocols 7, 500.

46. Kesimer, M., Ehre, C., Burns, K.A., Davis, C.W., Sheehan, J.K., and Pickles, R.J. (2013). Molecular organization of the mucins and glycocalyx underlying mucus transport over mucosal surfaces of the airways. Mucosal Immunol 6, 379–392.

47. Ali, M., Lillehoj, E.P., Park, Y., Kyo, Y., and Kim, K.C. (2011). Analysis of the proteome of human airway epithelial secretions. Proteome science 9, 1–10.

48. Reid, C.J., Gould, S., and Harris, A. (1997). Developmental expression of mucin genes in the human respiratory tract. American Journal of Respiratory Cell and Molecular Biology 17, 592–598.

49. Vsevolozhskaya, O.A., Shi, M., Hu, F., and Zaykin, D.V. (2020). DOT: Gene-set analysis by combining decorrelated association statistics. PLOS Computational Biology 16, e1007819.

50. Zhao, Y., and Sun, L. (2019). On set-based association tests: Insights from a regression using summary statistics. Canadian Journal of Statistics.

51. Derkach, A., Lawless, J.F., and Sun, L. (2014). Pooled Association Tests for Rare Genetic Variants: A Review and Some New Results. Statist Sci 29, 302–321.

52. Li, M.X., Gui, H.S., Kwan, J.S., and Sham, P.C. (2011). GATES: a rapid and powerful gene-based association test using extended Simes procedure. Am J Hum Genet 88, 283–293.

53. Lamparter, D., Marbach, D., Rueedi, R., Kutalik, Z., and Bergmann, S. (2016). Fast and Rigorous Computation of Gene and Pathway Scores from SNP-Based Summary Statistics. PLOS Computational Biology 12, e1004714.

