## Supplemental Information for "A flexible summary-based colocalization method with application to the mucin Cystic Fibrosis lung disease modifier locus"

### Supplemental Figures

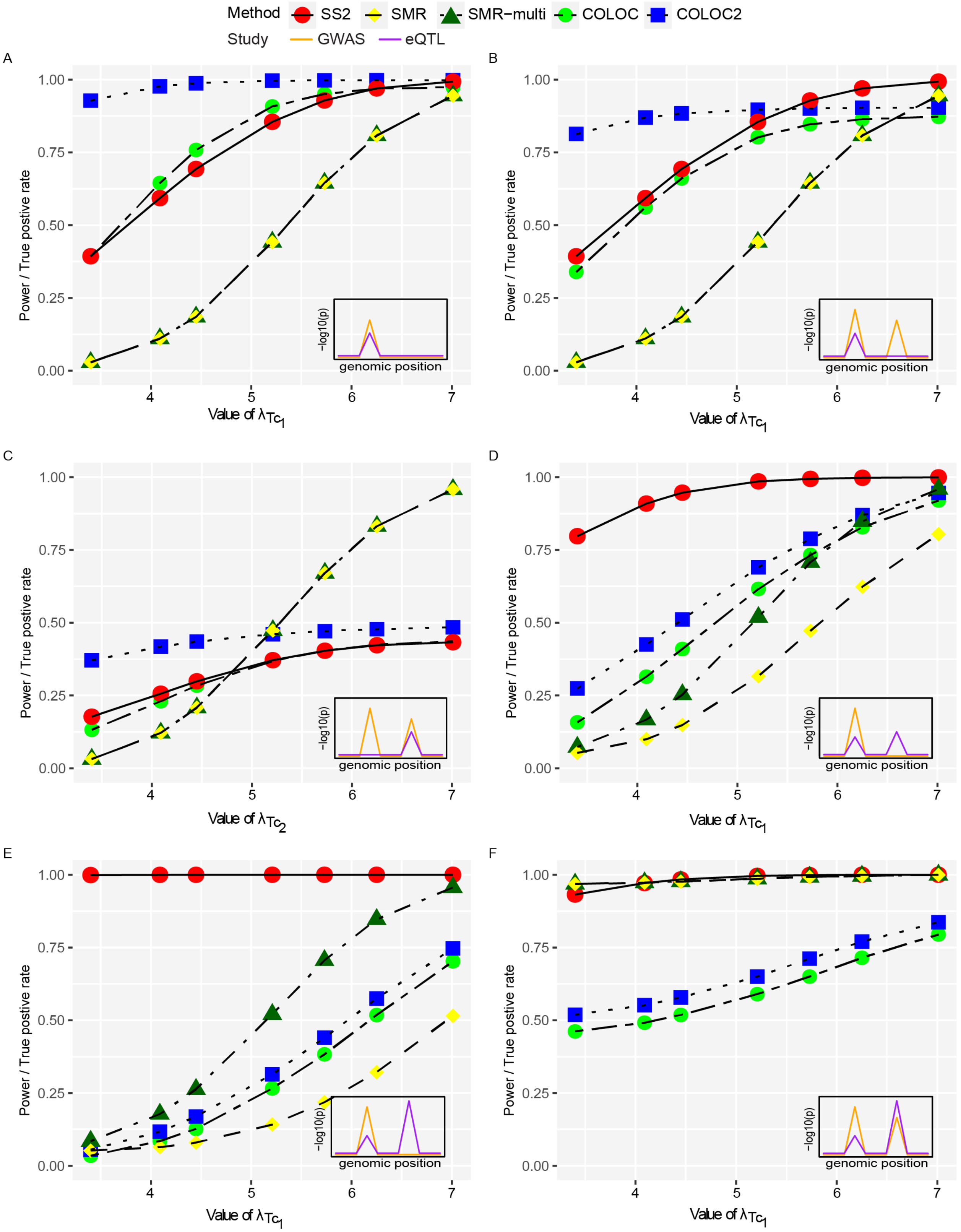

Method
SS2
SMR
SMR−multi
COLOC
COLOC2
Study
GWAS
eQTL
A
B
C
D
E
F
Power / True postive rate
−log10(p)
genomic position
Value of $\lambda_{Tc_1}$
Value of $\lambda_{Tc_2}$

**Figure S1. Empirical power of SS2, SMR, SMR-multi and true positive rates of COLOC and COLOC2 for testing a single gene for colocalization.** The LD pattern is simulated following the *SLC6A14* locus at chromosome X. For SS2, SMR[1] and SMR-multi,[2] the nominal type 1 error rate is set at alpha = 0.05. For COLOC[3] and COLOC2,[4] the false positive rates are controlled by applying the default 0.8 threshold (as recommended in [4]) for the colocalization posterior probability. In total, $10^4$ replications are simulated to obtain the empirical power and true positive rates. The x-axis represents parameter values for $\lambda_{T_{c1}}$ or $\lambda_{T_{c2}}$ (3.4, 4.09, 4.45, 5.21, 5.73 and 7.01) such that 0.01, 0.05, 0.1, 0.3, 0.5 and 0.9 power is achieved to detect the eQTL association at significance level $10^{-8}$. The subplot in the bottom right of each plot provides a general visualization of GWAS (orange line) and eQTL (purple line) colocalization patterns in a region of interest.

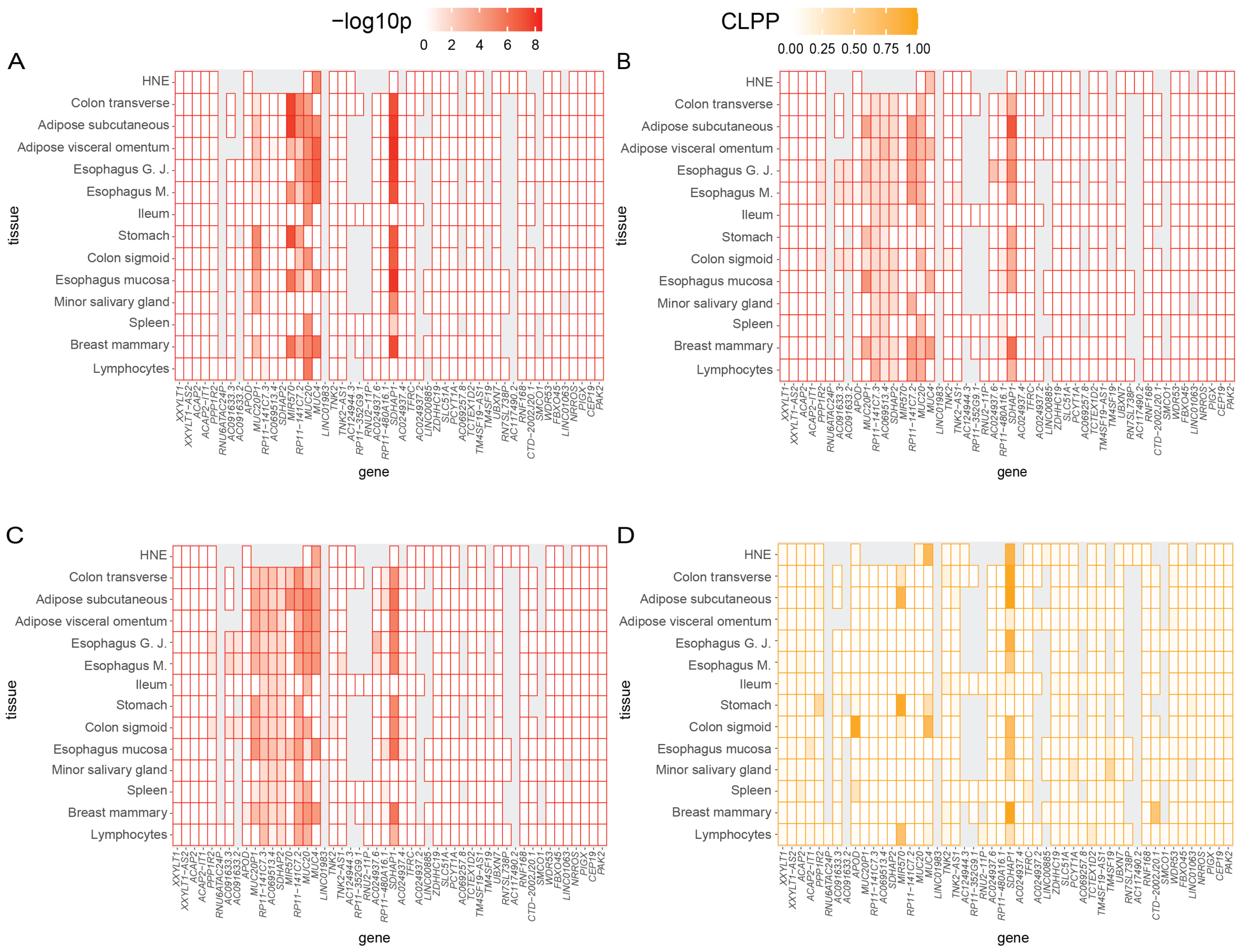


**Figure S2. Heatmaps of colocalization evidence across genes and tissues for a 1Mb region encompassing the peak lung-associated variants at chromosome 3: (A) SS2, (B) SMR, (C) SMR-multi, and (D) COLOC.** In each panel, each cell shows the colocalization evidence for the specified tissue and gene calculated from SNPs within 0.1Mb of the peak GWAS variant. The genes on the x-axis are within 1Mb on either side of the peak GWAS variant and are ordered by their chromosomal positions. Grey indicates insufficient expression levels attained for the gene in the tissue under study. (A): the color intensity corresponds to the SS2 colocalization evidence as measured by $-\log_{10}$(SS2 p-value), with red representing $-\log_{10}(p) = 8.5$ and white representing eQTL evidence for the corresponding gene and tissue does not pass the stage 1 test. (B) and (C): the color intensity corresponds to the SMR and SMR-multi colocalization evidence as measured by $-\log_{10}$(SMR p-value) and $-\log_{10}$(SMR-multi p-value), respectively; with red representing $-\log_{10}(p) = 8.5$ and white representing eQTL evidence for the corresponding gene and tissue does not pass the eQTL p-value threshold ($5\times10^{-8}$). (D): the color intensity corresponds to the COLOC colocalization evidence as measured by colocalization posterior probability (CLPP) ranging from 0 to 1. The eQTL analyses used for all gene/tissue pairs are those conducted by GTEx[5] version 8 release, except for the HNE eQTLs. eQTL analyses in HNE is conducted using FastQTL[6] with RNA-sequencing of HNE from 94 CF Canadians enrolled in the Canadian CF Gene Modifier study. Esophagus G.J represents Esophagus Gastroesophageal Junction; Esophagus M. represents Esophagus Muscularis; Ileum represents Small Intestine Terminal Ileum; Lymphocytes represents Cells EBV-transformed lymphocytes.

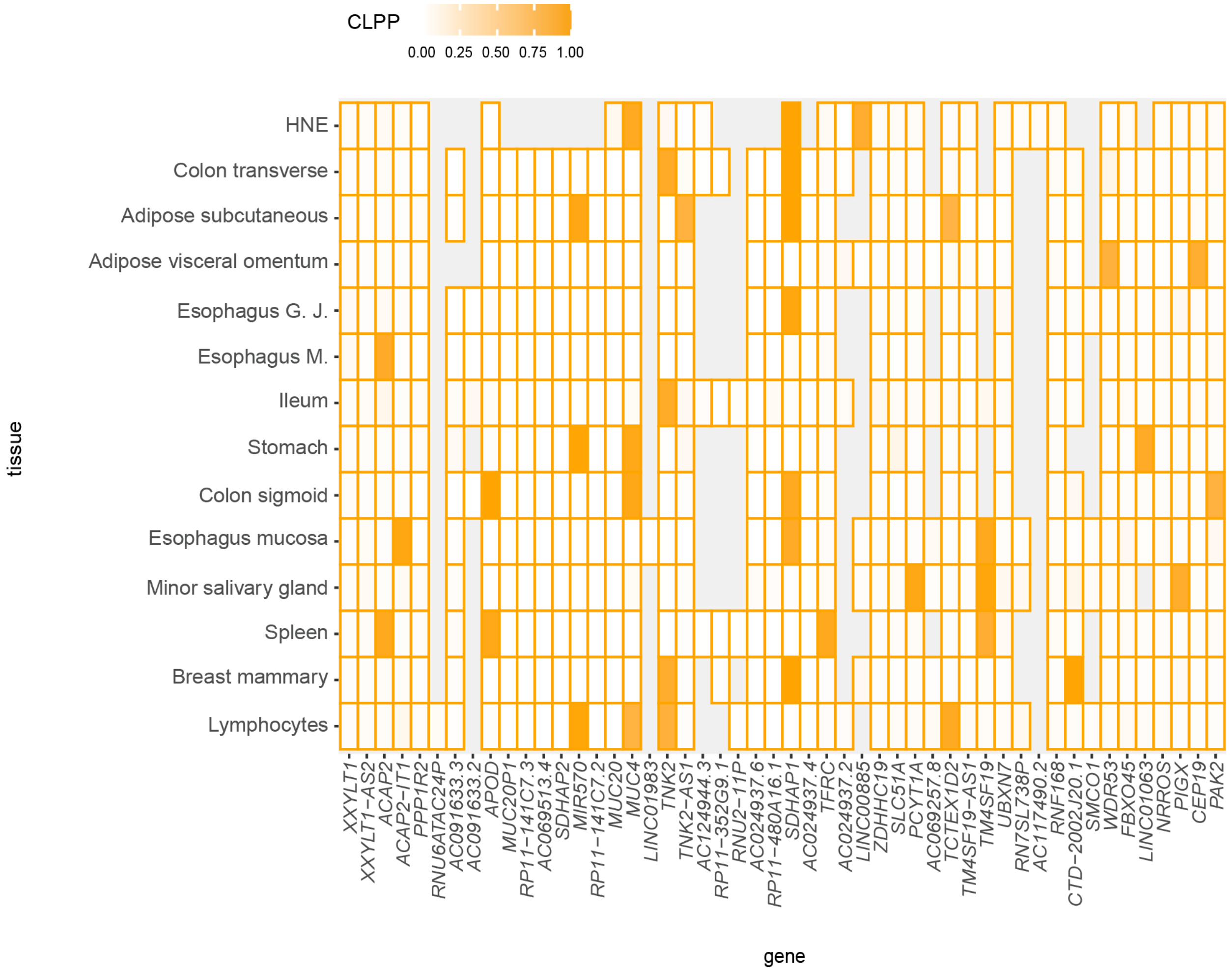


**Figure S3. Heatmap of colocalization evidence across genes and tissues for a 1Mb region encompassing the peak lung-associated variants at chromosome 3 by COLOC2.** COLOC2 analysis is conducted based on the likelihood from each single gene-by-tissue pair, calculated from SNPs within 0.1Mb of the peak GWAS variants. The color intensity corresponds to the COLOC2 colocalization evidence as measured by colocalization posterior probability (CLPP) ranging from 0 to 1. Grey indicates insufficient expression levels attained for the gene in the tissue under study. The eQTL data used for all gene/tissue pairs was sourced from GTEx version 8 release, with exception to the HNE eQTLs. eQTL analysis in HNE is conducted using FastQTL with RNA-sequencing of HNE from 94 CF Canadians enrolled in the Canadian CF Gene Modifier study. Esophagus G.J represents Esophagus Gastroesophageal Junction;

Esophagus M. represents Esophagus Muscularis; Ileum represents Small Intestine Terminal Ileum; Lymphocytes represents Cells EBV-transformed lymphocyte.

### Supplemental Tables

| Null Scenarios | $H_{01}$ | $H_{02}$ | $H_{03,(1)}$ | $H_{03,(2)}$ | $H_{04,(1)}$ | $H_{04,(2)}$ |
|---|---|---|---|---|---|---|
| Parameter Values | $\lambda_{Z_c} = 0$; $\lambda_{T_c} = 0$ | $\lambda_{Z_c} = 0$; $\lambda_{T_c} = 5.48$ | $\lambda_{Z_c} = 6.57$; $\lambda_{T_c} = 0$ | $\lambda_{Z_c} = 5.73$ $\lambda_{T_c} = 0$ | $\lambda_{Z_c} = 6.57$; $\lambda_{T_c} = 5.48$ | $\lambda_{Z_c} = 5.73$ $\lambda_{T_c} = 5.48$ |

**Table S1. Parameter settings under composite null hypothesis when conducting colocalization tests for a single gene with overlapping samples.** Each column corresponds to a specific null scenario when there is no colocalization. The values of $\lambda_{Zc}$ and $\lambda_{Tc}$ represent the standardized true effect size of a GWAS associated variant and an eQTL variant, respectively. For example, a GWAS association, $\lambda_{Zc}$ is set to be 0, 5.73, or 6.57 such that 0, 0.5 or 0.8 power is achieved to detect the signal at the significance level of $10^{-8}$. If there is an eQTL, $\lambda_{Tc}$ is set to be 5.48 such that 0.4 power is achieved to detect signal at the significance level of $10^{-8}$. $H_{01}$ represents the scenario when there are no SNP-phenotype associations and no eQTL; $H_{02}$ represents the scenario when there are no SNP-phenotype associations but eQTLs are present; $H_{03,(1)}$ and $H_{03,(2)}$ represent two scenarios where SNP-phenotype associations are present but no eQTL; $H_{04,(1)}$ and $H_{04,(2)}$ represent two scenarios where both SNP-phenotype association and eQTL are present, but occur at two independent SNPs. See Section *Simulation Details for Overlapping Samples* for other simulation details.

| Null Scenarios | Type I error of SS2 | | | | |
|---|---|---|---|---|---|
| | Correlation =0 | Correlation =0.3 | Correlation =0.5 | Correlation =0.7 | Correlation =0.9 |
| $H_{01}$ | 0.0033 | 0.0031 | 0.0033 | 0.0031 | 0.0030 |
| $H_{02}$ | 0.0430 | 0.0426 | 0.0425 | 0.0430 | 0.0425 |
| $H_{03,(1)}$ | 0.0243 | 0.0271 | 0.0266 | 0.0269 | 0.0259 |
| $H_{03,(2)}$ | 0.0178 | 0.0202 | 0.0199 | 0.0190 | 0.0202 |
| $H_{04,(1)}$ | 0.0257 | 0.0281 | 0.0277 | 0.0277 | 0.0269 |
| $H_{04,(2)}$ | 0.0187 | 0.0208 | 0.0206 | 0.0209 | 0.0217 |

**Table S2. Empirical type I error rates of SS2 when 50% of the participants in the eQTL study overlap with the GWAS study with varying phenotypic correlations.** The LD pattern at the simulated region follows that at the *MUC20/MUC4* locus. Simulations are performed under the null hypothesis of no colocalization and with varying phenotypic correlation (0.3, 0.5, 0.7 and 0.9). For comparison, the type I error rate when no samples are overlapping is also shown (correlation = 0). Table S1 defines the parameters used under each null scenario. The nominal type I error is set at alpha = 0.05. In total, $10^4$ replications are simulated for each null scenario. See Section *Simulation Details for Overlapping Samples* for other simulation details.

| Null Scenarios | Type I error of SS2 | | | |
|---|---|---|---|---|
| | correlation=0.3 | correlation=0.5 | correlation=0.7 | correlation=0.9 |
| $H_{01}$ | 0.0015 | 0.0024 | 0.0038 | 0.0039 |
| $H_{02}$ | 0.0373 | 0.0375 | 0.0385 | 0.0382 |
| $H_{03,(1)}$ | 0.0229 | 0.0242 | 0.0241 | 0.0224 |
| $H_{03,(2)}$ | 0.0241 | 0.0251 | 0.0254 | 0.0231 |
| $H_{04,(1)}$ | 0.0183 | 0.0181 | 0.0175 | 0.0174 |
| $H_{04,(2)}$ | 0.0188 | 0.0186 | 0.0176 | 0.0185 |

**Table S3. Empirical type I error rates of SS2 when 100% of the participants in the eQTL study overlap with the GWAS study, with varying phenotypic correlations.** The LD pattern at the simulated region follows that at the *MUC20/MUC4 locus.* Simulations are performed under the null hypothesis of no colocalization and with varying phenotypic correlation (0.3, 0.5, 0.7 and 0.9). Table S1 defines the parameters used under each null scenario. The nominal type I error is set at alpha = 0.05. In total, $10^4$ replications are simulated for each null scenario. See Section *Simulation Details for Overlapping Samples* for other simulation details.

| Null Scenarios | Type I error of SS2 | | | | |
|---|---|---|---|---|---|
| | eQTL sample size =200 | eQTL sample size =200 | eQTL sample size =300 | eQTL sample size =400 | eQTL sample size =500 |
| $H_{01}$ | 0.0037 | 0.0040 | 0.0053 | 0.0065 | 0.0086 |
| $H_{02}$ | 0.0495 | 0.0478 | 0.0487 | 0.0504 | 0.0493 |
| $H_{03,(1)}$ | 0.0266 | 0.0265 | 0.0227 | 0.0255 | 0.0243 |
| $H_{03,(2)}$ | 0.0288 | 0.0274 | 0.0233 | 0.0263 | 0.0255 |
| $H_{04,(1)}$ | 0.0199 | 0.0165 | 0.0222 | 0.0202 | 0.0261 |
| $H_{04,(2)}$ | 0.0190 | 0.0166 | 0.0218 | 0.0210 | 0.0255 |

**Table S4. Empirical type I error rates of SS2 when 100% of the participants in the eQTL study overlap with the GWAS study, with phenotypic correlation 0.5, and varying sample sizes of the eQTL study.** The LD pattern at the simulated region follows that at the *MUC20/MUC4* locus. Simulations are performed under the null hypothesis of no colocalization and with varying eQTL sample size (100, 200, 300, 400 or 500). Table S1 defines the parameters used under each null scenario. The nominal type I error is set at alpha = 0.05. In total, $10^4$ replications are simulated for each null scenario. See Section *Simulation Details for Overlapping Samples* for other simulation details.

| Null Scenarios | Type I error of SS2 | | | | |
|---|---|---|---|---|---|
| | eQTL sample size =100 | eQTL sample size =200 | eQTL sample size =300 | eQTL sample size =400 | eQTL sample size =500 |
| $H_{01}$ | 0.0037 | 0.009 | 0.0122 | 0.0177 | 0.0216 |
| $H_{02}$ | 0.0495 | 0.0502 | 0.0122 | 0.0572 | 0.0576 |
| $H_{03,(1)}$ | 0.0266 | 0.0258 | 0.0122 | 0.024 | 0.0252 |
| $H_{03,(2)}$ | 0.0288 | 0.0276 | 0.0122 | 0.0262 | 0.0262 |
| $H_{04,(1)}$ | 0.0199 | 0.0185 | 0.0122 | 0.0209 | 0.0296 |
| $H_{04,(2)}$ | 0.019 | 0.019 | 0.0122 | 0.0211 | 0.0304 |

**Table S5. Empirical type I error rates of SS2 when 100% of the participants in the eQTL study overlap with the GWAS study, with strong phenotypic correlation 0.9, and varying sample sizes of the eQTL study.** The LD pattern at the simulated region follows that at the *MUC20/MUC4* locus. Simulations are performed under the null hypothesis of no colocalization and with varying eQTL sample size (100, 200, 300, 400 or 500). Table S1 defines the parameters used under each null scenario. The nominal type I error is set at alpha = 0.05. In total, $10^4$ replications are simulated for each null scenario. See Section *Simulation Details for Overlapping Samples* for other simulation details.

| Null Scenarios | Parameter Values |
| --- | --- |
| $H_{01}$ | $\lambda_{Z_c} = 0$; $\lambda_{T_c} = 0$ |
| $H_{02}$ | $\lambda_{Z_c} = 0$; $\lambda_{T_c} = 7.01$ |
| $H_{03}$ | $\lambda_{Z_c} = 5.73$; $\lambda_{T_c} = 0$ |
| $H_{04}$ | $\lambda_{Z_c} = 5.73$; $\lambda_{T_c} = 7.01$ |

**Table S6. Parameter settings under composite null hypothesis when conducting colocalization tests for a single gene with independent samples.** Each row corresponds to a specific null scenario when there is no colocalization. $H_{01}$ represents the scenario when there is no SNP-phenotype association and no eQTL; $H_{02}$ represents the scenario when there is no SNP-phenotype association but eQTLs are present; $H_{03}$ represents the scenario where SNP-phenotype associations are present but no eQTL; $H_{04}$ represents the scenario where both SNP-phenotype association and eQTL are present, but occur at two independent SNPs. Values of $\lambda_{Z_c}$ and $\lambda_{T_c}$ represent the standardized true effect size of a GWAS associated variant and an eQTL variant, respectively. GWAS association, $\lambda_{Z_c}$ is set to be 5.73 such that 0.5 power is achieved to detect the signal at the significance level of $10^{-8}$. If there is an eQTL, $\lambda_{T_c}$ is set to be 7.01 such that 0.9 power is achieved to detect the signal at the significance level of $10^{-8}$. See Section *Simulation Details for a Single Gene-by-Tissue Pair* for other simulation details.

| Scenarios | Parameter Values |
|---|---|
| Scenario 1: one shared SNP for GWAS and eQTL study (Figure 2A and Figure S1A) | $\lambda_{Z_{c1}} = 6.57$; $\lambda_{Z_{c2}} = 0$<br>$\lambda_{T_{c1}} = 3.40, 4.45, 5.21, 5.73, 6.25$ or $7.01$;<br>$\lambda_{T_{c2}} = 0$ |
| Scenario 2: two independent GWAS SNPs, and one eQTL SNP is shared with the first GWAS SNP (Figure 2B and Figure S1B) | $\lambda_{Z_{c1}} = 6.57$; $\lambda_{Z_{c2}} = 5.73$<br>$\lambda_{T_{c1}} = 3.40, 4.45, 5.21, 5.73, 6.25$ or $7.01$;<br>$\lambda_{T_{c2}} = 0$ |
| Scenario 3: two independent GWAS SNPs, and one eQTL SNP is shared with the second GWAS SNP (Figure 2C and Figure S1C) | $\lambda_{Z_{c1}} = 6.57$; $\lambda_{Z_{c2}} = 5.73$<br>$\lambda_{T_{c1}} = 0$;<br>$\lambda_{T_{c2}} = 3.40, 4.45, 5.21, 5.73, 6.25$ or $7.01$ |
| Scenario 4: two independent eQTL SNPs, one GWAS SNP colocalizes with the eQTL SNP; the non-overlapping eQTL association is mild (Figure 2D and Figure S1D) | $\lambda_{Z_{c1}} = 6.57$; $\lambda_{Z_{c2}} = 0$<br>$\lambda_{T_{c1}} = 3.40, 4.45, 5.21, 5.73, 6.25$ or $7.01$;<br>$\lambda_{T_{c2}} = 5.73$ |
| Scenario 5: two independent eQTL SNPs, one GWAS SNP co-localizes with the eQTL SNP; the non-overlapping eQTL association is strong (Figure 2E and Figure S1E) | $\lambda_{Z_{c1}} = 6.57$; $\lambda_{Z_{c2}} = 0$<br>$\lambda_{T_{c1}} = 3.40, 4.45, 5.21, 5.73, 6.25$ or $7.01$;<br>$\lambda_{T_{c2}} = 7.01$ |

| | |
|---|---|
| Scenario 6: two independent GWAS SNPs, two independent eQTL SNPs and both SNPs are shared (Figure 2F and Figure S1F) | $\lambda_{Z_{c1}} = 6.57$; $\lambda_{Z_{c2}} = 5.73$<br>$\lambda_{T_{c1}} = 3.40, 4.45, 5.21, 5.73, 6.25$ or $7.01$;<br>$\lambda_{T_{c2}} = 7.01$ |

**Table S7. Parameter settings of the six alternative scenarios simulated to access the power/true positive rate of different methods.** $\lambda_{Z_{c_1}}$ and $\lambda_{Z_{c_2}}$ denote the standardized true effect sizes of two GWAS associated variants, while $\lambda_{T_{c_1}}$ and $\lambda_{T_{c_2}}$ denote the standardized true effect sizes of two eQTL variants. For the GWAS associated variants, $\lambda_{Z_{c_1}}$ is set to be 6.57 such that 0.8 power is achieved to detect the GWAS signal at the significance level $10^{-8}$; $\lambda_{Z_{c_2}}$ is set to be 0 or 5.73 such that 0 or 0.5 power is achieved to detect the GWAS signal at the significance level $10^{-8}$. For the eQTL variants, $\lambda_{T_{c_1}}$ is set to be 0, 3.4, 4.09, 4.45, 5.21, 5.73 or 7.01 such that 0, 0.01, 0.05, 0.1, 0.3, 0.5 or 0.9 power is achieved to detect the eQTL signal at the significance level $10^{-8}$; same for the value of $\lambda_{T_{c2}}$. See Section *Simulation Details for a Single Gene-by-Tissue Pair* for other simulation details.

| Locus | Proportion of genes with eQTL association but do not colocalize | FWER of SS2 | | | | |
|---|---|---|---|---|---|---|
| | | 100 genes | 200 genes | 300 genes | 400 genes | 500 genes |
| *MUC20/MUC4* | 0% | 0.0352 | 0.0348 | 0.0364 | 0.0367 | 0.0370 |
| | 20% | 0.0240 | 0.0272 | 0.0274 | 0.0283 | 0.0283 |
| | 40% | 0.0247 | 0.0267 | 0.0274 | 0.0291 | 0.0292 |
| | 60% | 0.0213 | 0.0227 | 0.0253 | 0.0279 | 0.0291 |
| | 80% | 0.0194 | 0.0223 | 0.0258 | 0.0293 | 0.0309 |
| | 100% | 0.0213 | 0.0270 | 0.0293 | 0.0311 | 0.0332 |
| *SLC6A14* | 0% | 0.0058 | 0.0055 | 0.0055 | 0.0053 | 0.0054 |
| | 20% | 0.0050 | 0.0031 | 0.0025 | 0.0020 | 0.0018 |
| | 40% | 0.0032 | 0.0021 | 0.0016 | 0.0014 | 0.0012 |
| | 60% | 0.0026 | 0.0017 | 0.0014 | 0.0012 | 0.0010 |
| | 80% | 0.0021 | 0.0014 | 0.0011 | 0.0010 | 0.0009 |
| | 100% | 0.0018 | 0.0012 | 0.0010 | 0.0009 | 0.0008 |

**Table S8. Empirical family-wise error rates (FWERs) of SS2 with different number of genes.** The LD pattern at the simulated region follows that at the *MUC20/MUC4* and *SLC6A14* loci. Simulations are conducted using a different number of genes (columns) and a varying proportion of genes with eQTL association (rows). The eQTL peaks are randomly generated from 6 different intervals (50%-60%, 60%-70%, 70%-80%, 80%-90%, 90%-95%, 95%-100% power is achieved to detect the eQTL association at the significance level of $10^{-8}$) with probabilities defined by the proportion of the -log10(maximum eQTL p-value) within each interval observed at the corresponding locus. The height of the GWAS peak is set at 5.06 on the -log10p scale such that 10% power is achieved to detect the GWAS association at the significance level of $10^{-8}$. None of the eQTL peaks colocalize with the GWAS peak for FWER evaluation. In total, $10^5$ replications are simulated to evaluate FWER of 0.05. The empirical FWER is calculated by counting the proportion of $10^5$ replications where at least one gene has a false colocalization claim. See Section *Simulation Details for Multiple Genes-Tissue Pairs* for other simulation details.

| Locus | Proportion of genes with eQTL association but do not colocalize | FWER of SMR | | | | |
|---|---|---|---|---|---|---|
| | | 100 genes | 200 genes | 300 genes | 400 genes | 500 genes |
| *MUC20/MUC4* | 0% | 0.0002 | 0.0004 | 0.0006 | 0.0007 | 0.0008 |
| | 20% | 0.0021 | 0.0011 | 0.0008 | 0.0006 | 0.0004 |
| | 40% | 0.0011 | 0.0005 | 0.0003 | 0.0002 | 0.0002 |
| | 60% | 0.0006 | 0.0003 | 0.0002 | 0.0002 | 0.0001 |
| | 80% | 0.0005 | 0.0003 | 0.0002 | 0.0001 | 0.0001 |
| | 100% | 0.0004 | 0.0002 | 0.0002 | $7.00\text{x}10^{-5}$ | $7.00\text{x}10^{-5}$ |
| *SLC6A14* | 0% | 0.0002 | 0.0003 | 0.0005 | 0.0005 | 0.0007 |
| | 20% | 0.0052 | 0.0023 | 0.0016 | 0.0012 | 0.0009 |
| | 40% | 0.0024 | 0.0012 | 0.0008 | 0.0006 | 0.0005 |
| | 60% | 0.0016 | 0.0008 | 0.0005 | 0.0004 | 0.0003 |
| | 80% | 0.0012 | 0.0006 | 0.0004 | 0.0002 | 0.0002 |
| | 100% | 0.0009 | 0.0004 | 0.0003 | 0.0002 | 0.0001 |

**Table S9. Empirical family-wise error rates (FWERs) of SMR with different number of genes.** The LD pattern at the simulated region follows that at the *MUC20/MUC4* and *SLC6A14* loci. SMR is conducted under the default setting such that a SNP is picked if only if the eQTL p-value is less than $5x10^{-8}$. Simulations are conducted using a different number of genes (columns) and a varying proportion of genes with eQTL association (rows). The eQTL peaks are randomly generated from 6 different intervals (50%-60%, 60%-70%, 70%-80%, 80%-90%, 90%-95%, 95%-100% power is achieved to detect the eQTL association at the significance level of $10^{-8}$) with probabilities defined by the proportion of the -log10(maximum eQTL p-value) within each interval observed at the corresponding locus. The height of the GWAS peak is set at 5.06 on the -log10p scale such that 10% power is achieved to detect the GWAS association at the significance level of $10^{-8}$. None of the eQTL peaks colocalize with the GWAS peak for FWER evaluation. In total, $10^5$ replications are simulated to evaluate FWER of 0.05. The empirical FWER is calculated by counting the proportion of $10^5$ replications where at least one gene has a false colocalization claim. See Section *Simulation Details for Multiple Genes-Tissue Pairs* for other simulation details.

| Locus | Proportion of genes with eQTL association but do not colocalize | FWER of SMR-multi | | | | |
|---|---|---|---|---|---|---|
| | | 100 genes | 200 genes | 300 genes | 400 genes | 500 genes |
| *MUC20/MUC4* | 0% | 0.0002 | 0.0004 | 0.0006 | 0.0007 | 0.0008 |
| | 20% | 0.0015 | 0.0006 | 0.0004 | 0.0003 | 0.0002 |
| | 40% | 0.0006 | 0.0003 | 0.0002 | 0.0001 | 0.0001 |
| | 60% | 0.0003 | 0.0002 | 0.0001 | $9.00 \times 10^{-5}$ | $7.00 \times 10^{-5}$ |
| | 80% | 0.0002 | 0.0002 | $8.00 \times 10^{-5}$ | $6.00 \times 10^{-5}$ | $6.00 \times 10^{-5}$ |
| | 100% | 0.0003 | 0.0002 | $7.00 \times 10^{-5}$ | $6.00 \times 10^{-5}$ | $5.00 \times 10^{-5}$ |
| *SLC6A14* | 0% | 0.0002 | 0.0004 | 0.0005 | 0.0005 | 0.0007 |
| | 20% | 0.0035 | 0.0015 | 0.0009 | 0.0007 | 0.0005 |
| | 40% | 0.0014 | 0.0007 | 0.0004 | 0.0003 | 0.0002 |
| | 60% | 0.0009 | 0.0004 | 0.0002 | 0.0002 | 0.0002 |
| | 80% | 0.0007 | 0.0003 | 0.0002 | 0.0002 | 0.0002 |
| | 100% | 0.0004 | 0.0002 | 0.0002 | 0.0001 | 0.0001 |

**Table S10. Empirical family-wise error rates (FWERs) of Multi-SNP-based SMR test (SMR-multi) with different number of genes.** The LD pattern at the simulated region follows that at the *MUC20/MUC4* and *SLC6A14* loci. Multi-SNP-based SMR test

(SMR-multi) is conducted under the default setting such that a SNP is picked if only if the eQTL p-value is less than $5x10^{-8}$. Simulations are conducted using a different number of genes (columns) and a varying proportion of genes with eQTL association (rows). The eQTL peaks are randomly generated from 6 different intervals (50%-60%, 60%-70%, 70%-80%, 80%-90%, 90%-95%, 95%-100% power is achieved to detect the eQTL association at the significance level of $10^{-8}$) with probabilities defined by the proportion of the -log10(maximum eQTL p-value) within each interval observed at the corresponding locus. The height of the GWAS peak is set at 5.06 on the -log10p scale such that 10% power is achieved to detect the GWAS association at the significance level of $10^{-8}$. None of the eQTL peaks colocalize with the GWAS peak for FWER evaluation. In total, $10^5$ replications are simulated to evaluate FWER of 0.05. The empirical FWER is calculated by counting the proportion of $10^5$ replications where at least one gene has a false colocalization claim. See Section *Simulation Details for Multiple Genes-Tissue Pairs* for other simulation details.

| Locus | Proportion of genes with eQTL association but do not colocalize | False positive rate of COLOC | | | | |
|---|---|---|---|---|---|---|
| | | 100 genes | 200 genes | 300 genes | 400 genes | 500 genes |
| *MUC20/MUC4* | 0% | 0.0265 | 0.0489 | 0.0704 | 0.0912 | 0.1118 |
| | 20% | 0.0223 | 0.0406 | 0.0587 | 0.0758 | 0.0927 |
| | 40% | 0.0180 | 0.0320 | 0.0460 | 0.0597 | 0.0729 |
| | 60% | 0.0134 | 0.0232 | 0.0330 | 0.0424 | 0.0515 |
| | 80% | 0.0088 | 0.0142 | 0.0197 | 0.0252 | 0.0300 |
| | 100% | 0.0009 | 0.0017 | 0.0022 | 0.0030 | 0.0037 |
| *SLC6A14* | 0% | 0.0350 | 0.0634 | 0.0902 | 0.1157 | 0.1400 |
| | 20% | 0.0269 | 0.0509 | 0.0735 | 0.0952 | 0.1162 |
| | 40% | 0.0218 | 0.0408 | 0.0592 | 0.0766 | 0.0938 |
| | 60% | 0.0163 | 0.0304 | 0.0441 | 0.0574 | 0.0703 |
| | 80% | 0.0111 | 0.0201 | 0.0289 | 0.0373 | 0.0458 |
| | 100% | 0.0057 | 0.0093 | 0.0128 | 0.0164 | 0.0198 |

**Table S11. Empirical false positive rates of COLOC with different number of genes.** The LD pattern at the simulated region follows that at the *MUC20/MUC4* and *SLC6A14* loci. Simulations are conducted using a different number of genes (columns) and a varying proportion of genes with eQTL association (rows). The eQTL peaks are randomly generated from 6 different intervals (50%-60%, 60%-70%, 70%-80%, 80%-90%, 90%-95%, 95%-100% power is achieved to detect the eQTL association at the significance level of $10^{-8}$) with probabilities defined by the proportion of the -log10(maximum eQTL p-value) within each interval observed at the corresponding locus. The height of the GWAS peak is set at 5.06 on the -log10p scale such that 10% power is achieved to detect the GWAS association at the significance level of $10^{-8}$. None of the eQTL peaks colocalize with the GWAS peak for FWER evaluation. In total, $10^5$ replications are simulated to evaluate the false positive rates by applying the 0.8 threshold (as recommended by [4]) for the colocalization posterior probability. The empirical false positive rate for COLOC is calculated by counting the proportion of $10^5$ replications where at least one gene has a false colocalization claim. See Section *Simulation Details for Multiple Genes-Tissue Pairs* for other simulation details.

| Locus | Proportion of genes with eQTL association but do not colocalize | False positive rate of COLOC2 | | | | |
|---|---|---|---|---|---|---|
| | | 100 genes | 200 genes | 300 genes | 400 genes | 500 genes |
| *MUC20/MUC4* | 0% | 0.0020 | 0.0034 | 0.0048 | 0.0061 | 0.0073 |
| | 20% | 0.0023 | 0.0023 | 0.0023 | 0.0023 | 0.0023 |
| | 40% | 0.0025 | 0.0026 | 0.0027 | 0.0027 | 0.0027 |
| | 60% | 0.0033 | 0.0033 | 0.0033 | 0.0034 | 0.0034 |
| | 80% | 0.0038 | 0.0040 | 0.0040 | 0.0040 | 0.0040 |
| | 100% | 0.0044 | 0.0044 | 0.0044 | 0.0044 | 0.0044 |
| *SLC6A14* | 0% | $2.00 \times 10^{-5}$ | $5.00 \times 10^{-5}$ | $9.00 \times 10^{-5}$ | 0.0001 | 0.0002 |
| | 20% | 0.0046 | 0.0047 | 0.0047 | 0.0048 | 0.0049 |
| | 40% | 0.0054 | 0.0055 | 0.0055 | 0.0056 | 0.0056 |
| | 60% | 0.0068 | 0.0069 | 0.0070 | 0.0070 | 0.0070 |
| | 80% | 0.0079 | 0.0079 | 0.0080 | 0.0080 | 0.0080 |
| | 100% | 0.0088 | 0.0088 | 0.0089 | 0.0089 | 0.0089 |

**Table S12. Empirical false positive rates of COLOC2 with different number of genes.** The LD pattern at the simulated region follows that at the *MUC20/MUC4* and *SLC6A14* loci. Simulations are conducted using a different number of genes (columns) and a varying proportion of genes with eQTL association (rows). The eQTL peaks are randomly generated from 6 different intervals (50%-60%, 60%-70%, 70%-80%, 80%-90%, 90%-95%, 95%-100% power is achieved to detect the eQTL association at the significance level of $10^{-8}$) with probabilities defined by the proportion of the $-\log10$(maximum eQTL p-value) within each interval observed at the corresponding locus. The height of the GWAS peak is set at 5.06 on the $-\log10p$ scale such that 10% power is achieved to detect the GWAS association at the significance level of $10^{-8}$. None of the eQTL peaks colocalize with the GWAS peak for FWER evaluation. In total, $10^5$ replications are simulated to evaluate the false positive rates by applying the 0.8 threshold (as recommended by [4]) for the colocalization posterior probability. The empirical false positive rates for COLOC2 are calculated by counting the proportions of $10^5$ replications where at least one gene has a false colocalization claim. See Section *Simulation Details for Multiple Genes-Tissue Pairs* for other simulation details.

| Locus | eQTL height for the 5% of genes that colocalize | Power of SS2 | | | | |
|---|---|---|---|---|---|---|
| | | 100 genes | 200 genes | 300 genes | 400 genes | 500 genes |
| *MUC20/MUC4* | 5.48-5.73 | 0.8880 | 0.8668 | 0.8508 | 0.8379 | 0.8268 |
| | 5.73-5.98 | 0.8891 | 0.8671 | 0.8512 | 0.8381 | 0.8269 |
| | 5.98-6.25 | 0.8893 | 0.8672 | 0.8508 | 0.8376 | 0.8261 |
| | 6.25-6.57 | 0.8891 | 0.8667 | 0.8499 | 0.8368 | 0.8255 |
| | 6.57-7.01 | 0.8885 | 0.8658 | 0.8488 | 0.8352 | 0.8235 |
| *SLC6A14* | 5.48-5.73 | 0.7126 | 0.7104 | 0.7018 | 0.6938 | 0.6855 |
| | 5.73-5.98 | 0.7450 | 0.7337 | 0.7200 | 0.7090 | 0.6992 |
| | 5.98-6.25 | 0.7642 | 0.7468 | 0.7306 | 0.7183 | 0.7076 |
| | 6.25-6.57 | 0.7752 | 0.7547 | 0.7376 | 0.7247 | 0.7138 |
| | 6.57-7.01 | 0.7819 | 0.7601 | 0.7424 | 0.7295 | 0.7184 |

**Table S13. Power of SS2 with different number of genes.** The LD pattern at the simulated region follows that at the *MUC20/MUC4* and *SLC6A14* loci. The height of the GWAS peak is set at 5.06 on the -log10p scale such that 10% power is achieved to detect the GWAS association at the significance level of $10^{-8}$. Simulations are conducted using a different number of genes (columns) and a varying range of the eQTL height for the 5%

of genes that colocalize (rows). The eQTL peaks are set with 5 different intervals such that 40%-50%, 50%-60%, 60%-70%, 70%-80%, 80%-90% power is achieved to detect the eQTL association at the significance level of $10^{-8}$. The 95% of genes that do not colocalize are simulated under a mixture of $H_{03}$ and $H_{04}$, and details are demonstrated in the Supplemental Methods. The definitions of $H_{03}$ and $H_{04}$ are provided in Table S6. In total, $10^5$ replications are simulated to evaluate power at 0.05 significance level. The power is calculated by counting the proportion of $10^5$ replications where at least one gene is correctly identified with colocalization. See Section *Simulation Details for Multiple Genes-Tissue Pairs* for other simulation details.

| Locus | eQTL height for the 5% of genes that colocalize | Power of SMR | | | | |
|---|---|---|---|---|---|---|
| | | 100 genes | 200 genes | 300 genes | 400 genes | 500 genes |
| *MUC20/MUC4* | 5.48-5.73 | 0.8299 | 0.7907 | 0.7540 | 0.7257 | 0.7015 |
| | 5.73-5.98 | 0.8467 | 0.7993 | 0.7633 | 0.7359 | 0.7126 |
| | 5.98-6.25 | 0.8561 | 0.8072 | 0.7717 | 0.7448 | 0.7225 |
| | 6.25-6.57 | 0.8633 | 0.8149 | 0.7808 | 0.7542 | 0.7322 |
| | 6.57-7.01 | 0.8704 | 0.8234 | 0.7906 | 0.7646 | 0.7435 |
| *SLC6A14* | 5.48-5.73 | 0.8005 | 0.7486 | 0.7067 | 0.6749 | 0.6519 |
| | 5.73-5.98 | 0.8184 | 0.7599 | 0.7186 | 0.6879 | 0.6652 |
| | 5.98-6.25 | 0.8292 | 0.7699 | 0.7293 | 0.6995 | 0.6767 |
| | 6.25-6.57 | 0.8379 | 0.7796 | 0.7398 | 0.7102 | 0.6886 |
| | 6.57-7.01 | 0.8449 | 0.7886 | 0.7504 | 0.7213 | 0.7005 |

**Table S14. Power of SMR with different number of genes.** The LD pattern at the simulated region follows that at the *MUC20/MUC4* and *SLC6A14* loci. SMR is conducted under the default setting such that a SNP is picked if only if the eQTL p-value is less than $5x10^{-8}$. The height of the GWAS peak is set at 5.06 on the -log10p scale such that 10% power is achieved to detect the GWAS association at the significance level of

$10^{-8}$. Simulations are conducted using a different number of genes (columns) and a varying range of the eQTL height for the 5% of genes that colocalize (rows). The eQTL peaks are set with 5 different intervals such that 40%-50%, 50%-60%, 60%-70%, 70%-80%, 80%-90% power is achieved to detect the eQTL association at the significance level of $10^{-8}$. The 95% of genes that do not colocalize are simulated under a mixture of $H_{03}$ and $H_{04}$, and details are demonstrated in the Supplemental Methods. The definitions of $H_{03}$ and $H_{04}$ are provided in Table S6. In total, $10^5$ replications are simulated to evaluate power at 0.05 significance level. The power is calculated by counting the proportion of $10^5$ replications where at least one gene has correctly identified with colocalization. See Section *Simulation Details for Multiple Genes-Tissue Pairs* for other simulation details.

| Locus | eQTL height for the 5% of genes that colocalize | Power of SMR-multi | | | | |
|---|---|---|---|---|---|---|
| | | 100 genes | 200 genes | 300 genes | 400 genes | 500 genes |
| *MUC20/MUC4* | 5.48-5.73 | 0.7955 | 0.7432 | 0.6964 | 0.6615 | 0.6316 |
| | 5.73-5.98 | 0.8077 | 0.7477 | 0.7009 | 0.6655 | 0.6363 |
| | 5.98-6.25 | 0.8133 | 0.7510 | 0.7050 | 0.6702 | 0.6410 |
| | 6.25-6.57 | 0.8162 | 0.7536 | 0.7074 | 0.6734 | 0.6453 |
| | 6.57-7.01 | 0.8161 | 0.7542 | 0.7094 | 0.6758 | 0.6481 |
| *SLC6A14* | 5.48-5.73 | 0.7659 | 0.7003 | 0.6483 | 0.6077 | 0.5785 |
| | 5.73-5.98 | 0.7789 | 0.7068 | 0.6548 | 0.6155 | 0.5859 |
| | 5.98-6.25 | 0.7850 | 0.7118 | 0.6604 | 0.6215 | 0.5923 |
| | 6.25-6.57 | 0.7883 | 0.7154 | 0.6652 | 0.6266 | 0.5983 |
| | 6.57-7.01 | 0.7884 | 0.7166 | 0.6686 | 0.6311 | 0.6036 |

**Table S15. Power of Multi-SNP-based SMR test (SMR-multi) with different number of genes.** The LD pattern at the simulated region follows that at the *MUC20/MUC4* and *SLC6A14* loci. SMR-multi is conducted under the default setting such that a SNP is picked if only if the eQTL p-value is less than $5x10^{-8}$. The height of the GWAS peak is set at 5.06 on the -log10p scale such that 10% power is achieved to detect the GWAS association at the significance level of $10^{-8}$. Simulations are conducted using a different number of genes (columns) and a varying range of the eQTL height for the 5% of genes that colocalize (rows). The eQTL peaks are set with 5 different intervals such that 40%-50%, 50%-60%, 60%-70%, 70%-80%, 80%-90% power is achieved to detect the eQTL association at the significance level of $10^{-8}$. The 95% of genes that do not colocalize are simulated under a mixture of $H_{03}$ and $H_{04}$, and details are demonstrated in the Supplemental Methods. The definitions of $H_{03}$ and $H_{04}$ are provided in Table S6. In total, $10^5$ replications are simulated to evaluate power at 0.05 significance level. The power is calculated by counting the proportion of $10^5$ replications where at least one gene is correctly identified with colocalization. See Section *Simulation Details for Multiple Genes-Tissue Pairs* for other simulation details.

| Locus | eQTL height for the 5% of genes that colocalize | True positive rate of COLOC2 | | | | |
|---|---|---|---|---|---|---|
| | | 100 genes | 200 genes | 300 genes | 400 genes | 500 genes |
| *MUC20/MUC4* | 5.48-5.73 | 0.4976 | 0.5516 | 0.5783 | 0.5968 | 0.6105 |
| | 5.73-5.98 | 0.5193 | 0.5690 | 0.5942 | 0.6118 | 0.6249 |
| | 5.98-6.25 | 0.5353 | 0.5817 | 0.6051 | 0.6219 | 0.6343 |
| | 6.25-6.57 | 0.5477 | 0.5908 | 0.6129 | 0.6287 | 0.6405 |
| | 6.57-7.01 | 0.5581 | 0.5981 | 0.6184 | 0.6333 | 0.6439 |
| *SLC6A14* | 5.48-5.73 | 0.5801 | 0.6258 | 0.6407 | 0.6494 | 0.6555 |
| | 5.73-5.98 | 0.5979 | 0.6364 | 0.6497 | 0.6581 | 0.6638 |
| | 5.98-6.25 | 0.6118 | 0.6443 | 0.6565 | 0.6643 | 0.6697 |
| | 6.25-6.57 | 0.6232 | 0.6516 | 0.6625 | 0.6693 | 0.6745 |
| | 6.57-7.01 | 0.6330 | 0.6572 | 0.6669 | 0.6734 | 0.6781 |

**Table S16. True positive rates of COLOC2 with different number of genes.** The LD pattern at the simulated region follows that at the *MUC20/MUC4* and *SLC6A14* loci. The height of the GWAS peak is set at 5.06 on the -log10p scale such that 10% power is achieved to detect the GWAS association at the significance level of $10^{-8}$. Simulations are conducted using a different number of genes (columns) and a varying range of the eQTL height for the 5% of genes that colocalize (rows). The eQTL peaks are set with 5 different intervals such that 40%-50%, 50%-60%, 60%-70%, 70%-80%, 80%-90% power is achieved to detect the eQTL association at the significance level of $10^{-8}$. The 95% of genes that do not colocalize are simulated under a mixture of $H_{03}$ and $H_{04}$, and details are demonstrated in the Supplemental Methods. The definitions of $H_{03}$ and $H_{04}$ are provided in Table S6. In total, $10^5$ replications are simulated to evaluate the true positive rates by applying the 0.8 threshold (as recommended by [4]) for the colocalization posterior probability. The true positive rates for COLOC2 are calculated by counting the proportion of $10^5$ replications where at least one gene is correctly identified with colocalization. See Section *Simulation Details for Multiple Genes-Tissue Pairs* for other simulation details.

| Null Scenarios | False positive rate of COLOC2 | | | |
|---|---|---|---|---|
| | eQTL sample size = 200 | eQTL sample size = 500 | eQTL sample size = 1000 | eQTL sample size = 2000 |
| $H_{01}$ | 0.0022 | 0.0025 | 0.0024 | 0.0021 |
| $H_{02}$ | 0.0468 | 0.0468 | 0.0467 | 0.0467 |
| $H_{03}$ | 0.0880 | 0.0692 | 0.0550 | 0.0417 |
| $H_{04}$ | 0.0012 | 0.0012 | 0.0012 | 0.0012 |

**Table S17. False positive rates of COLOC2 for a single hypothesis test with different sample sizes for the eQTL study.** The LD pattern at the simulated region follows that at the *SLC6A14* locus. Sample size is fixed at 2000 participants in the GWAS study, and is varied for the eQTL study (columns). Each row corresponds to a specific null scenario when there is no colocalization. See Table S6 for the explanations of each null scenario with the corresponding parameter settings. The false positive rates are calculated by applying the 0.8 threshold (as recommended by [4]) for the colocalization posterior probability. In total, $10^4$ replications are simulated for each null scenario. See Section *Simulation Details for a Single Gene-by-Tissue Pair* for other simulation details.

| Null Scenarios | Type I error of SMR | | Type I error of SMR-multi | |
|---|---|---|---|---|
| | (eQTL p<0.01) | (eQTL p<0.05) | (eQTL p<0.01) | (eQTL p<0.05) |
| $H_{01}$ | 0.0067 | 0.0071 | 0.003 | $6.00 \times 10^{-4}$ |
| $H_{02}$ | 0.0392 | 0.0392 | 0.0162 | 0.006 |
| $H_{03}$ | 0.0766 | 0.0934 | 0.0843 | 0.0807 |
| $H_{04}$ | 0.0404 | 0.0404 | 0.0483 | 0.0606 |

**Table S18. Empirical type I error rates of SMR and Multi-SNP-based SMR test (SMR-multi) for a single hypothesis test under eQTL p-value thresholds 0.01 and 0.05.** SMR and Multi-SNP-based SMR test (SMR-multi) are conducted under the setting such that a SNP is picked if only if the eQTL p-value is less than 0.01(eQTL p<0.01) or 0.05 (eQTL p<0.05). The LD pattern at the simulated region follows that at the *MUC20/MUC4* locus. Each row corresponds to a specific null scenario when there is no colocalization. Table S6 defines the parameters used under each null scenario. The nominal type I error is set at alpha = 0.05. In total, $10^4$ replications are simulated for each null scenario.

| Null Scenarios | Type I error of SS2 | | | | |
|---|---|---|---|---|---|
| | Correlation =0 | Correlation =0.3 | Correlation =0.5 | Correlation =0.7 | Correlation =0.9 |
| $H_{01}$ | 0.0033 | 0.0091 | 0.0249 | 0.0422 | 0.0489 |
| $H_{02}$ | 0.0430 | 0.052 | 0.0643 | 0.0884 | 0.1404 |
| $H_{03,(1)}$ | 0.0243 | 0.0222 | 0.0254 | 0.0282 | 0.0320 |
| $H_{03,(2)}$ | 0.0178 | 0.0300 | 0.0338 | 0.0405 | 0.0501 |
| $H_{04,(1)}$ | 0.0257 | 0.0226 | 0.0258 | 0.0288 | 0.0317 |
| $H_{04,(2)}$ | 0.0187 | 0.0312 | 0.0336 | 0.0397 | 0.0478 |

**Table S19: Empirical type I error rates of SS2 when all of 2000 participants in the eQTL study overlap with the GWAS study, with varying phenotypic correlations.** The LD pattern at the simulated region follows that at the *MUC20/MUC4* locus. Simulations are performed under the null hypothesis of no colocalization and with varying phenotypic correlation (0.3, 0.5, 0.7 and 0.9). For comparison, the type I error rate when no samples are overlapping is also shown (correlation = 0). Table S1 defines the parameters used under each null scenario. The nominal type I error is set at alpha = 0.05. In total, $10^4$ replications are simulated for each null scenario. See Section *Simulation Details for Overlapping Samples* for other simulation details.

### Supplemental Methods

#### Simulation Details for a Single Gene-by-Tissue Pair

We generate the GWAS summary statistics $\boldsymbol{Z}$ and the eQTL summary statistics $\boldsymbol{T}$ from $N(\Sigma\Lambda_Z, \Sigma)$ and $N(\Sigma\Lambda_T, \Sigma)$, respectively, where $\Sigma$ is the LD matrix from the locus of interest. $\Lambda_z = \left(\lambda_{Z_1}, \lambda_{Z_2}, \dots \lambda_{Z_c}, \dots \lambda_{Z_m}\right)^{\top}$ and $\Lambda_T = \left(\lambda_{T_1}, \lambda_{T_2}, \dots \lambda_{T_c}, \dots \lambda_{T_m}\right)^{\top}$ are vectors with each component being the true standardized effect size of the corresponding SNP, respectively for GWAS and eQTL analyses. In particular, $\lambda_{Z_c}$ and $\lambda_{T_c}$ represent the value for the associated SNP from the GWAS and eQTL studies, respectively. When there is SNP-phenotype association, we use the lead GWAS SNP from our CF study[7,8] as the associated SNP with value $\lambda_{Z_c}$, and we define a SNP ($r < 0.002$ and $> -0.002$ with the associated) SNP as an independent eQTL SNP with value $\lambda_{T_c}$.

For type I error evaluation, we consider all four scenarios of the composite null hypothesis. For example, under $H_{01}$ when there is no SNP-phenotype association and no eQTL, $\lambda_{Z_c} = 0$ and $\lambda_{T_c} = 0$. In contrast, under $H_{04}$ when both SNP-phenotype association and eQTL are present but occurring at independent SNPs, $\lambda_{Z_c} \neq 0$ and $\lambda_{T_c} \neq 0$, and we set $\lambda_{Z_c} = 5.73$ and $\lambda_{T_c} = 7.01$ to be consistent with the previous simulation study in [7]. Table S6 provides details for the parameter settings under the four null scenarios. Under each scenario, $10^4$ replications are simulated.

To study power, we simulate six scenarios. For scenarios with two association peaks, we used the lead GWAS SNP from our CF study at the locus to be the first associated SNP, while the second associated SNP is defined as the next adjacent SNP with $r < 0.002$ and $> -0.002$ with the first associated SNP. We vary the magnitude of eQTL evidence to measure the relationship between power and colocalization strength. Table S7 provides details for the parameter settings under the six alternative scenarios. Under each scenario, $10^4$ replications are simulated.

#### Simulation Details for Multiple Genes-Tissue Pairs

The GWAS summary statistics are generated from $N(\Sigma\Lambda_Z, \Sigma)$ with $\lambda_{Z_c} = 4.5$, and $\Sigma$ is simulated based on the LD within the *MUC20/MUC4* and *SLC6A14* loci. The two loci are defined by including SNPs within 0.1Mb of either side of the top associated GWAS SNP from the CF study. For FWER evaluation with one GWAS association at the simulated locus, we generate 600 sets of independent eQTL summary statistics corresponding to eQTL analyses of 600 genes of interest under $H_{03}$ or $H_{04}$, where different proportions of genes with eQTL evidence under $H_{04}$ are considered (0%, 20%, 40%, 60%, 80% and 100%). For a given gene with eQTL signals under $H_{04}$, the eQTL summary statistics are generated from $N(\Sigma\Lambda_T, \Sigma)$ where the value of $\lambda_{T_c}$ is randomly generated from 6 different intervals (50%-60%, 60%-70%, 70%-80%, 80%-90%, 90%-95%, 95%-100% power is achieved to detect the eQTL association at the significance level of $10^{-8}$) with probabilities according to the proportion of the -log10 (maximum eQTL p-value) within each interval observed at the locus. For other genes with eQTL under $H_{03}$, the eQTL summary statistics are generated from $\mathrm{N}(0, \Sigma)$.

For power evaluation, with the same GWAS peak, we set 5% of genes with eQTL signals as colocalizing with the GWAS signal (Scenario 1 in Table 1). For the *SLC6A14* locus, we set 47.5% of genes as having no eQTL signal (under $H_{03}$), and 47.5% of genes as having eQTL signals that are independent from the GWAS signal (under $H_{04}$); while modeling the *MUC20/MUC4* locus, we set 19% of genes under $H_{03}$, and 76% of genes under $H_{04}$. The GWAS associated SNP and the eQTL associated SNPs are selected in the same way in section *Simulation Details for a Single Gene-by-Tissue Pair*. For genes under the alternative, the eQTL summary statistics are generated from $\mathrm{N}(\Sigma\Lambda_T, \Sigma)$, where the value of $\lambda_{T_c}$ is randomly selected from an interval (i.e. [5.48, 5.73]). For genes with the eQTL under $H_{04}$, $\lambda_{T_c}$ is selected from 4.45 to 9.48 such that 10% to 100% power is achieved at genome-wide significance level $10^{-8}$. We evaluate power as the average increase in strength of eQTL among the 5% of genes under the alternative, and consider 5 different intervals: [5.48, 5.73], [5.73, 5.98], [5.98, 6.25], [6.25, 6.57] and [6.57, 7.01]. Under each scenario, $10^5$ replications are simulated.

#### Simulation Details for Overlapping Samples

Given the sample size $n_{GWAS}$ and $n_{eQTL}$ for a GWAS and eQTL study, respectively, we randomly select $n_{GWAS}$ individuals from the genotype data of the CF lung GWAS study.[8] A region is defined which included the SNPs 0.1Mb around the top GWAS SNP at the *MUC4/MUC20* locus. All or half of the participants in the eQTL study are randomly selected and simulated such that they are overlapping with the participants in the GWAS.

For individual $i$, let $g_{1i}$ and $g_{2i}$ denote the genotype of the associated SNP for the GWAS and eQTL study respectively. We simulate a pair of phenotypes for each individual $\binom{y_{1i}}{y_{2i}} = \binom{g_{1i}\beta_1+\epsilon_{1i}}{g_{2i}\beta_2+\epsilon_{2i}}$, where $\beta_1$ and $\beta_2$ denote the effect sizes for the GWAS and eQTL studies, respectively. Hormozdiari et al[9] has shown that

$$\begin{aligned} \lambda_{Z_c} &= \frac{\beta_1}{\sqrt{\text{var}\,(\epsilon_{1i})}} \times \sqrt{n_{GWAS}}; \\ \lambda_{T_c} &= \frac{\beta_2}{\sqrt{\text{var}\,(\epsilon_{2i})}} \times \sqrt{n_{eQTL}}. \end{aligned} \tag{1}$$

We set different values for $\lambda_{Z_c}$ and $\lambda_{T_c}$ under the composite null hypothesis, and then calculate the corresponding values for $\beta_1$ and $\beta_2$ according to equation (1). For example, when the GWAS signal and eQTL signal are distinct $(H_{04})$, we set $\Lambda_{T_c} = 5.48$ to mimic the observed eQTL peak for *MUC4* in HNE. Given 100 participants in the eQTL study $(n_{eQTL} = 100)$ and $var(\epsilon_{2i}) = 1$, we obtain $\beta_1$= 0.123. We consider two different values of $\lambda_{Z_c}$ (5.73 or 6.57) such that 0.5 or 0.8 power is achieved to detect the GWAS association at the genome-wide significance level of $10^{-8}$. Given 2000 participants in the GWAS study and $var(\epsilon_{2i}) = 1$, we obtain $\beta_1$= 0.128 and 0.147, respectively. Parameter settings for $\lambda_{Z_c}$ and $\lambda_{T_c}$ under the composite null hypothesis is provided in Table S1. $\binom{\epsilon_{1i}}{\epsilon_{2i}}$ ~MVN ($(\binom{0}{0}, \Sigma_\epsilon)$), where $\Sigma_\epsilon$ is the covariance matrix with the diagonal elements set to be 1. If an individual is included in both studies, the off-diagonal terms are set to C which represents the phenotypic correlation; otherwise it is set to be 0. Simulation studies are conducted for different values of phenotypic correlation, C= 0.3, 0.5, 0.7 or 0.9. We perform univariate linear regression to obtain the marginal summary statistics and then apply SS2. Under each scenario, $10^4$ replications are simulated.

#### Covariance of Summary Statistics from Meta-analyses with Related Individuals

Suppose the GWAS meta-analysis consists of $C$ sub-studies with sample sizes $n_c, c = 1,2, \dots C$. Let $\hat{\beta}_c = (\hat{\beta}_{c,1}, \dots, \hat{\beta}_{c,m})^\top$ and $Z_c = (Z_{c,1}, Z_{c,2}, \cdots, Z_{c,m})^\top$ denote the marginal effect sizes and Z-scores, respectively, from study c. Let $Z_{\text{meta}} = (Z_{\text{meta},1}, Z_{\text{meta},2}, \dots Z_{\text{meta},m})^\top$, where $Z_{\text{meta},j}$ represents the Z-score obtained from the meta-analysis for SNP $j$. Then

$$Z_{meta,j} = \frac{\sum_{c=1}^{C} w_{c,j}\hat{\beta}_{c,j}}{\sqrt{\sum_{c=1}^{C} w_{c,j}}} \quad (2)$$

$$= \frac{\sum_{c=1}^{C} \sqrt{w_{c,j}} Z_{c,j}}{\sqrt{\sum_{c=1}^{C} w_{c,j}}}, \quad (3)$$

where $w_{c,j}$ represents the weight for study c and can take different forms (i.e. the inverse variance of $\hat{\beta}_{c,j}$). In particular, for the sample size weighted meta-analysis, $w_{c,j} = n_c$ and $Z_{meta,j} = \frac{\sum_{c=1}^{C}\sqrt{n_c} z_{c,j}}{\sqrt{n\lambda_c}}$, where $\lambda_c$ is the genomic control.[10] For SNP $j$ and SNP $l$,

$$\mathrm{cov}\left(Z_{meta,j}, Z_{meta,l}\right) = \mathrm{cov}\left(\frac{\sum_{c=1}^{C}\sqrt{w_{c,j}} Z_{c,j}}{\sqrt{\sum_{c=1}^{C} w_{c,j}}}, \frac{\sum_{c=1}^{C}\sqrt{w_{c,l}} Z_{c,l}}{\sqrt{\sum_{c=1}^{C} w_{c,l}}}\right)$$

$$= \frac{\sum_{c=1}^{C}\sqrt{w_{c,j}}\sqrt{w_{c,l}}}{\sqrt{\sum_{c=1}^{C} w_{c,j}}\sqrt{\sum_{c=1}^{C} w_{c,l}}} \mathrm{cov}\left(Z_{c,j}, Z_{c,l}\right). \quad (4)$$

Therefore, the covariance of meta-Z scores between two SNPs is a weighted sum of covariance measures for the sub-studies.

If all participants in the sub-study c are independent, we consider fitting the following single-marker model with additional covariates to obtain the marginal summary statistics:

$$Y = G_j\beta + X\alpha + \epsilon \text{ with } \epsilon \sim N(0, \sigma^2 I), \quad (5)$$

for SNP $j. Y = \left(y_1, \ldots, y_{n_c}\right)^{\top}$ denote a $n_c \times 1$ vector of phenotypic values, where each component $y_i$ is the phenotype for the $i$th individual. Assume there are m SNPs at the locus, $G_j = \left(G_{j1}, \dots, G_{jn_c}\right)^\top$ is a $n_c \times 1$ vector of genotype for SNP $j$,where each component $G_{ij}$ is the genotype for the $i$th individual. $X$ is a $n_c \times q$ matrix of covariates (i.e. sex and age) and $\alpha$ is a $q \times 1$ vector of fixed effects for the covariates, including the intercept.

We define $U_j = \left(G_j, X\right)$, which is a $n_c \times (q+1)$ matrix that contains the genotype for SNP $j$ and additional covariates. Let $B = (1, 0, 0, \dots, 0)$, a $1 \times (1+q)$ vector with the first component to be 1 and other components to be 0. We obtain $\hat{\beta}_{c,j}$ and $Z_{c,j}$ by the least square method:

$$\hat{\beta}_{c,j} = B\left(U_j^\top U_j\right)^{-1} U_j^\top Y \quad ; \tag{6}$$

$$Z_{c,j} = \frac{B\left(U_j^\top U_j\right)^{-1} U_j^\top Y}{\sqrt{\hat{\sigma}_j^2 B\left(U_j^\top U_j\right)^{-1} B^\top}}, \tag{7}$$

where $\hat{\sigma}_j^2$ is the maximum likelihood estimator (MLE) of $\sigma_j$ in model (5). For SNP $j$ and SNP $l$,

$$\begin{aligned} \operatorname{cov}\left(Z_{c,j}, Z_{c,l}\right) &= \operatorname{cov}\left(\frac{B\left(U_j^\top U_j\right)^{-1} U_j^\top Y}{\sqrt{\hat{\sigma}_j^2 B\left(U_j^\top U_j\right)^{-1} B^\top}}, \frac{B(U_l^\top U_l)^{-1} U_l^\top Y}{\sqrt{\hat{\sigma}_l^2 B(U_l^\top U_l)^{-1} B^\top}}\right) \\ &= \frac{B\left(U_j^\top U_j\right)^{-1} U_j^\top}{\sqrt{\hat{\sigma}_j^2 B\left(U_j^\top U_j\right)^{-1} B^\top}} \sigma^2 I \frac{U_l (U_l^\top U_l)^{-1} B^\top}{\sqrt{\hat{\sigma}_l^2 B(U_l^\top U_l)^{-1} B^\top}}, \end{aligned}$$

where $\hat{\sigma}_i^2$ and $\hat{\sigma}_l^2$ approaches $\sigma^2$ in probability. By Slutsky's theorem, the asymptotically covariance is

$$\operatorname{cov}(Z_{c,j}, Z_{c,l}) = \frac{B\left(U_j^\top U_j\right)^{-1} U_j^\top U_l (U_l^\top U_l)^{-1} B^\top}{\sqrt{B\left(U_j^\top U_j\right)^{-1} B^\top B (U_l^\top U_l)^{-1} B^\top}}. \tag{8}$$

Zheng et al[11] showed equation (8) can be simplified such that

$$\begin{aligned} \operatorname{cov}\left(Z_{c,j}, Z_{c,l}\right) &= \frac{G_j^\top (I - P_X) G_l}{\sqrt{G_j^t (I - P_X) G_j} \sqrt{G_l^t (I - P_X) G_l}} \\ &= \operatorname{cor}\left(\hat{\epsilon}_{G_j|X}, \hat{\epsilon}_{G_l|X}\right), \end{aligned} \tag{9}$$

where $P_X = X(X^\top X)^{-1} X^\top . \hat{\epsilon}_{G_j|X}$ denotes the residuals obtained by regressing $G_j$ on $X$, for $j = 1, .. m$. In particular, if both phenotypes and genotypes are standardized, and there are no additional covariates expect the genotypes $(B = 1$ and $U_j = G_j)$, equation (9) can be simplified as

$$\operatorname{cov}(Z_{c,j}, Z_{c,l}) = \frac{G_j^\top G_l}{\sqrt{G_j^\top G_j G_l^\top G_l}}.$$

The asymptotic covariance of Z scores between two SNPs is their pairwise Pearson correlation coefficient.[11,12]

If the sub-study $c$ contains related samples, the first approach to obtain the summary statistics (for SNP $j$) is considering a linear fixed-effect model:

$$Y = G_j \beta + X\alpha + \epsilon, \text{ with } \epsilon \sim N(0, \Sigma_\epsilon), \tag{10}$$

where $\Sigma_\epsilon$ denote the matrix that contains information about the sample relatedness (i.e. the kinship coefficient matrix). If $\Sigma_\epsilon$ is known, let $H$ denote the Cholesky decomposition of $\Sigma_\epsilon^{-1}$ such that $H^t H = \Sigma_\epsilon^{-1}$. We define $Y^* = HY, X^* = HX, G_j^* =$

$HG_j$, $U_j^* = HU_j$ and $\epsilon^* = H\epsilon$. By the generalized least square (GLS) method, the effect size and Z-score for SNP $j$ is

$$\begin{aligned}\hat{\beta}_{c,j} &= B\left(U_j^{*\top}U_j^*\right)^{-1}U_j^{*\top}Y^* \\ &= B\left(U_j^\top\Sigma_\epsilon^{-1}U_j\right)^{-1}U_j^\top\Sigma_\epsilon^{-1}Y; \qquad (11)\\ Z_{c,j} &= \frac{B\left(U_j^{*\top}U_j^*\right)^{-1}U_j^*Y^*}{\sqrt{B\left(U_j^{*\top}U_j^*\right)^{-1}B^\top}} \\ &= \frac{B\left(U_j^\top\Sigma_\epsilon^{-1}U_j\right)^{-1}U_j^\top\Sigma_\epsilon^{-1}Y}{\sqrt{B\left(U_j^\top\Sigma_\epsilon^{-1}U_j\right)^{-1}B^\top}}. \qquad (12)\end{aligned}$$

Similar to equation (9),

$$\begin{aligned}\mathrm{cov}\left(Z_{c,j}, Z_{c,l}\right) &= \frac{G_j^{*\top}(I-P_{X^*})G_l^*}{\sqrt{G_j^*(I-P_{X^*})G_j^*}\sqrt{G_l^{*\top}(I-P_{X^*})G_l^*}}, \text{ with } P_{X^*} = X^*(X^{*\top}X^*)^{-1}X^{*\top}; \\ &= \frac{G_j^\top P^* G_l}{\sqrt{G_j^\top P^* G_j}\sqrt{G_l^\top P^* G_l}}, \text{ with } P^* = \Sigma_\epsilon^{-1} - \Sigma_\epsilon^{-1}X(X^\top\Sigma_\epsilon^{-1}X)^{-1}X^\top\Sigma_\epsilon^{-1}. \quad (13)\end{aligned}$$

If $\Sigma_\epsilon$ is unknown, we can get a consistent estimator of $\Sigma_\epsilon$ first,[13] say $\hat{\Sigma}_\epsilon$, and then obtain the effect size and Z-score by using $\hat{\Sigma}_\epsilon$:

$$\hat{\beta}_{c,j}\left(\hat{\Sigma}_\epsilon\right) = B\left(U_j^\top\hat{\Sigma}_\epsilon^{-1}U_j\right)^{-1}U_j^\top\hat{\Sigma}_\epsilon^{-1}Y; \qquad (14)$$

$$Z_{c,j}\left(\hat{\Sigma}_\epsilon\right) = \frac{B\left(U_j^\top\hat{\Sigma}_\epsilon^{-1}U_j\right)^{-1}U_j^\top\hat{\Sigma}_\epsilon^{-1}Y}{\sqrt{B\left(U_j^\top\hat{\Sigma}_\epsilon^{-1}U_j\right)^{-1}B^\top}}. \qquad (15)$$

By Slutsky theorem, equation (13) is the asymptotic covariance of Z-scores. In practice, we estimate the covariance by $\hat{\Sigma}_\epsilon$:

$$\mathrm{cov}\left(Z_{c,j}, Z_{c,l}\right) = \frac{G_j^\top\hat{P}^*G_l}{\sqrt{G_j^\top\hat{P}^*G_j}\sqrt{G_l^\top\hat{P}^*G_l}}, \text{ with } \hat{P}^* = \hat{\Sigma}_\epsilon^{-1} - \hat{\Sigma}_\epsilon^{-1}X\left(X^\top\hat{\Sigma}_\epsilon^{-1}X\right)^{-1}X^\top\hat{\Sigma}_\epsilon^{-1}. \quad (16)$$

Secondly, the linear mixed-effect model (LMM) with a random intercept is another commonly used approach for related samples:[14,15]

$$Y = G_j\beta + X\alpha + Q + \epsilon, \text{ with } Q \sim N\left(0, \sum_{k=1}^{K} \tau_k \Phi_k\right) \text{ and } \epsilon \sim N(0, \sigma^2 I). \quad (17)$$

$Q$ is the random intercept; $\Phi_k$ is a known genetic relationship matrix (i.e. the kinship coefficient matrix times 2), and $\tau_k$ is the corresponding variance component parameter. $Q$ and $\epsilon$ are assumed to be independent. Given the observed covariates $X$ and $G_j$ ($j = 1, \ldots, m$), the distribution of $Y$ is $N\left(G_j\beta + X\gamma, \Sigma_\epsilon\right)$, where $\Sigma_\epsilon = \sum_{k=1}^{K} \tau_k \Phi_k + \sigma^2 I$.

When $\Sigma_\epsilon$ is known, the effect size in equation (11) is MLE based on the marginal distribution of $Y$ and is also the minimum variance unbiased estimate.[16] Therefore, we can estimate the covariance of $Z$ -scores by equation (13) under this case, which is the same as the covariance derived under linear fixed-effect model in (10).

When $\Sigma_\epsilon$ is unknown, the variance parameters contained in $\Sigma_\epsilon$ can be estimated by EM algorithm for maximum likelihood (ML) estimates or restricted maximum likelihood (REML) estimates.[16,17] Then the effect size is calculated by equation (14),[16] and this procedure has been implemented in the R package nlme.[18] Therefore, the Z-scores and its covariance can be calculated by equation (15) and equation (16), respectively, using variance component estimates from MLE or REML. Other types of summary statistics such as the score-based statistics have been implemented in R packages GMMAT based on the LMM assumption, with the covariance estimated by AI-REML algorithm.[19]

#### Type I Error Rate Control for a Single Hypothesis Testing

Let $H_0$ denote the null hypothesis that there is no colocalization given a specific gene-by-tissue pair, including multiple different scenarios: $H_{01}, H_{02}, H_{03}$, and $H_{04}$. The definitions of $H_{01}, H_{02}, H_{03}$, and $H_{04}$ are provided in Table S6. For a specific gene-by-tissue pair, let $R_1$ and $R_2$ denote the event that the stage 1 eQTL test and the stage 2 SS test, respectively, provide significant p-values. Let $R_{SS2}$ denote the event where the SS2 test rejects the null hypothesis of no colocalization. Thus, $R_{SS2} = R_1 \cap R_2$. The type I error rate for testing a single gene-by-tissue pair is

$$
\begin{aligned}
P(R_{SS2} \mid H_0) &= P(R_2 \mid R_1, H_0) \times P(R_1 \mid H_0) \\
&= \sum_{i=1}^{4} P(R_{SS2} \mid H_{0i}) \times P(H_{0i} \mid H_0) \\
&= \sum_{i=1}^{4} P(R_2 \mid R_1, H_{0i}) \times P(R_1 \mid H_{0i}) \times P(H_{0i} \mid H_0) \qquad (18) \\
&= \sum_{i=1}^{4} P(R_2 \mid H_{0i}) \times P(R_1 \mid H_{0i}) \times P(H_{0i} \mid H_0) \qquad (19) \\
&\leq \sum_{i=1}^{4} \alpha \times P(H_{0i} \mid H_0) \qquad (20) \\
&= \alpha
\end{aligned}
$$

From equation (18) to equation (19), $P(R_2 \mid H_{0i}) = P(R_2 \mid R_1, H_{0i})$ because the stage 2 test is independent of the stage 1 test. We access the distribution of the stage 2 test statistics by conditioning on the observed eQTL summary statistics, where the variation of the eQTL summary statistics is only considered in the stage 1 test. In equation (20), for $i = 1$ or 3 when there is no eQTL, the stage 1 test controls the type I error rate within $\alpha$

and thus $P(R_1 \mid H_{0i}) \leq \alpha;$ for $i = 2$ or 4, the stage 2 test controls the type I error rate within $\alpha$ and thus $P(R_2 \mid H_{0i}) \leq \alpha$. Therefore, $P(R_{SS2} \mid H_0) \leq \alpha$.

#### Upper Bound of Family-wise Error Rate for Multiple Hypothesis Testing

Suppose we have a locus of interest, and we are interested in testing colocalization for $K$ gene-by-tissue pairs at this locus. The following proof formally derives the upper bound of the Family-wise error rate (FWERs) when the $K$ tests consist of different scenarios. Let $\alpha$ denote the nominal significance level for the SS2 test. Let $H_{0,K}$ denote the null hypothesis that there are no colocalizations for the $K$ gene-by-tissue pairs at the locus. If there is no SNP-phenotype association at the locus (denoted by $H_{0,K,nsp}$), the $K$ tests are under $H_{01}$ or $H_{02}$; if there is SNP-phenotype association at the locus (denoted by $H_{0,K,sp}$), the $K$ tests are under $H_{03}$ or $H_{04}$. Let $R_{SS2}^*$ denote the event that the SS2 test rejects at least one tests among the $K$ tests. Therefore. for a locus with $K$ tests, the FWER of the SS2 test is

$$
\begin{aligned}
&P\left(R_{SS2}^* \mid H_{0,K}\right) \\
&= P\left(R_{SS2}^* \mid H_{0,K,nsp}\right) \times P\left(H_{0,K,nsp} \mid H_{0,K}\right) \qquad (21) \\
&+P\left(R_{SS2}^* \mid H_{0,K,sp}\right) \times P\left(H_{0,K,sp} \mid H_{0,K}\right). \qquad (22)
\end{aligned}
$$

$P\left(H_{0,K,sp} \mid H_{0,K}\right)$ represents the probability that the locus has phenotype-SNP association given there are no colocalizations for the $K$ gene-by-tissue pairs, which is unknown in practice. Therefore, we need to find the maximum values for both $P\left(R_{SS2}^* \mid H_{0,K,nsp}\right)$ in (21) and $P\left(R_{SS2}^* \mid H_{0,K,sp}\right)$ in (22) to obtain the upper bound for $P\left(R_{SS2}^* \mid H_{0,K}\right)$. Let $R_{1,P}$ represent the event that the stage 1 test has $P$ significant gene-by-tissue pairs. Let $R_{2,P}^*$ represents the event that the stage 2 test rejects at least one gene-by-tissue pairs among the $P$ significant gene-by-tissue pairs from the stage 1 test.

For a locus with no phenotype-SNP association (under $H_{0,K,nsp}$), the $K$ tests consist of mixture of $H_{01}$ or $H_{02}$. If there are $P$ significant gene-by-tissue pairs from the stage 1 test, the stage 2 test is implemented at significant level $\frac{\alpha}{P}$ per test by Bonferroni correction, and thus $P\left(R_{2,P}^* \mid R_{1,P}, H_{0,K,nsp}\right) \leq \frac{\alpha}{P} \times P = \alpha$. Therefore,

$$\begin{aligned} P\left(R_{SS2}^* \mid H_{0,K,nsp}\right) &= \sum_{P=1}^{K} P\left(R_{2,P}^* \mid R_{1,P}, H_{0,K,nsp}\right) \times P\left(R_{1,P} \mid H_{0,K,nsp}\right) \\ &\leq \sum_{P=1}^{K} \left(\frac{\alpha}{P} \times P\right) \times P\left(R_{1,P} \mid H_{0,K,nsp}\right) \\ &= \alpha \times \sum_{P=1}^{K} P\left(R_{1,P} \mid H_{0,K,nsp}\right) \\ &\leq \alpha \end{aligned} \qquad (23)$$

For a locus with phenotype-SNP association (under $H_{0,K,sp}$), we need to consider three cases separately: the $K$ tests are all under $H_{04}$; the $K$ tests are all under $H_{03}$; the $K$ tests consist of a mixture of $H_{03}$ and $H_{04}$. Let $H_{0,K,h4}, H_{0,K,h3}$ and $H_{0,K,h34}$ denote the three cases. respectively. Therefore,

$$\begin{aligned} P\left(R_{SS2}^* \mid H_{0,K,sp}\right) \quad &= P\left(R_{SS2}^* \mid H_{0,K,h4}\right) \times P\left(H_{0,K,h4} \mid H_{0,K,sp}\right) \\ &+ P\left(R_{SS2}^* \mid H_{0,K,h3}\right) \times P\left(H_{0,K,h3} \mid H_{0,K,sp}\right) \\ &+ P\left(R_{SS2}^* \mid H_{0,K,h34}\right) \times P\left(H_{0,K,h34} \mid H_{0,K,sp}\right). \end{aligned} \qquad (24)$$

Under $H_{0,K,h4}$, when all the $K$ tests are under $H_{04}$, $P\left(R_{2,P}^{*} \mid R_{1,P}, H_{0,K,nsp}\right) \leq \frac{\alpha}{P} \times P = \alpha$. Thus,

$$\begin{aligned} P\left(R_{SS2}^{*} \mid H_{0,K,h4}\right) &= \sum_{P=1}^{K} P\left(R_{2,P}^{*} \mid R_{1,P}, H_{0,K,h4}\right) \times P\left(R_{1,P} \mid H_{0,K,h4}\right) \\ &\leq \sum_{P=1}^{K} (\frac{\alpha}{P} \times P) \times P\left(R_{1,P} \mid H_{0,K,h4}\right) \\ &= \alpha \times \sum_{P=1}^{K} P\left(R_{1,P} \mid H_{0,K,h4}\right) \\ &\leq \alpha. \end{aligned} \tag{25}$$

Under $H_{0.K,h3}$ when all the $K$ tests are under $H_{03}$, the stage 1 test for each gene-by-tissue pair is implemented under the null that there is no eQTL, and adjust the $\alpha$ for the total number of tests by Bonferroni correction, $\frac{\alpha}{K}$. Thus,

$$\begin{aligned} P\left(R_{SS2}^{*} \mid H_{0,K,h3}\right) &\leq P\left(\text{ the stage 1 test rejects at least one gene-by-tissue pairs } \mid H_{0,K,h3}\right) \\ &\leq \frac{\alpha}{K} \times K \\ &= \alpha. \end{aligned} \tag{26}$$

So far we have proved that when all the $K$ tests are under $H_{03}$ or $H_{04}$, the upper bound on the FWER of the two-stage SS2 test is $\alpha$.

Under $H_{0,K,h34}$ when $K$ tests consist of $H_{03}$ and $H_{04}$, let $H_{0,K_1,K_2}$ denote the event that there are $K_1$ tests under $H_{04}$ and $K_2$ tests under $H_{03}$. We define $R_{1,P_1,P_2}$ to be the event that there are $P_1$ significant stage 1 eQTL tests among those $K_1$ tests and $P_2$ significant stage 1 eQTL tests among those $K_2$ tests. In this case,

$$P\left(R_{SS2}^* \mid H_{0,K,h_{34}}\right) = \sum_{K_1 \geq 1, K_2 \geq 1} P\left(R_{SS2}^* \mid H_{0,K_1,K_2}\right) \times P\left(H_{0,K_1,K_2} \mid H_{0,K,h_{34}}\right),$$

$$\begin{aligned}
P\left(R_{SS2}^* \mid H_{0,K_1,K_2}\right) = & \sum_{P_1 \geq 1, P_2 = 0} P\left(R_{2,p}^* \mid R_{1,P_1,P_2}, H_{0,K_1,K_2}\right) \times P\left(R_{1,P_1,P_2} \mid H_{0,K_1,K_2}\right) \\
& + \sum_{P_1 = 0, P_2 \geq 1} P\left(R_{2,p}^* \mid R_{1,P_1,P_2}, H_{0,K_1,K_2}\right) \times P\left(R_{1,P_1,P_2} \mid H_{0,K_1,K_2}\right) \\
& + \sum_{P_1 \geq 1, P_2 \geq 1} P\left(R_{2,p}^* \mid R_{1,P_1,P_2}, H_{0,K_1,K_2}\right) \times P\left(R_{1,P_1,P_2} \mid H_{0,K_1,K_2}\right) \\
& = \sum_{P_1=1}^{K_1} P\left(R_{2,p_1}^* \mid R_{1,P_1,0}, H_{0,K_1,K_2}\right) \times P\left(R_{1,P_1,0} \mid H_{0,K_1,K_2}\right) \qquad (27) \\
& + \sum_{P_2=1}^{K_2} P\left(R_{2,p_2}^* \mid R_{1,0,P_2}, H_{0,K_1,K_2}\right) \times P\left(R_{1,0,P_2} \mid H_{0,K_1,K_2}\right) \qquad (28) \\
& + \sum_{P_1 \geq 1, P_2 \geq 1} P\left(R_{2,p}^* \mid R_{1,P_1,P_2}, H_{0,K_1,K_2}\right) \times P\left(R_{1,P_1,P_2} \mid H_{0,K_1,K_2}\right). \qquad (29)
\end{aligned}$$

The term (27) demonstrates the scenario when all the significant stage 1 eQTL tests are within the $K_1$ gene-by-tissue pairs under $H_{04}$. In this scenario, $P = P_1$ and the stage 2 test is implemented at significance level $\frac{\alpha}{P_1}$ per test. Therefore, term (27)

$$\begin{aligned}
& \sum_{P_1=1}^{K_1} P\left(R_{2,p_1}^* \mid R_{1,P_1,0}, H_{0,K_1,K_2}\right) \times P\left(R_{1,P_1,0} \mid H_{0,K_1,K_2}\right) \\
& \leq \left(\frac{\alpha}{P_1} \times P_1\right) \times \sum_{P_1=1}^{K_1} P\left(R_{1,P_1,0} \mid H_{0,K_1,K_2}\right) \\
& = \alpha \times \sum_{P_1=1}^{K_1} P\left(R_{1,P_1,0} \mid H_{0,K_1,K_2}\right) \\
& = \alpha \times S_{(1,0)}, \qquad (30)
\end{aligned}$$

Where $S_{(1,0)} = \sum_{P_1=1}^{K_1} P\left(R_{1,P_1,0} \mid H_{0,K_1,K_2}\right)$ which represents the probability that there is at least one significant stage 1 eQTL test among the $K_1$ gene-by-tissue pairs but no significant stage 1 eQTL test among the $K_2$ gene-by-tissue pairs.

Let $\alpha^*$ denote the probability that the stage 2 test rejects at least one gene-by-tissue pairs for those $P$ significant stage 1 eQTL tests. The term $(28)$ demonstrates the scenario when all the significant stage 1 eQTL tests are within the $K_2$ gene-by-tissue pairs under $H_{03}$. In this scenario, $P = P_2$; the stage 2 test is implemented at significance level $\frac{\alpha}{P_2}$ per test and $\alpha^* = P\left(R^*_{2,p_2} \mid R_{1,0,P_2}, H_{0,K_1,K_2}\right)$.

$$\begin{aligned}
&\sum_{P_2=1}^{K_2} P\left(R^*_{2,p_2} \mid R_{1,0,P_2}, H_{0,K_1,K_2}\right) \times P\left(R_{1,0,P_2} \mid H_{0,K_1,K_2}\right) \\
&= \alpha^* \times \sum_{P_2=1}^{K_2} P\left(R_{1,0,P_2} \mid H_{0,K_1,K_2}\right) \\
&= \alpha^* \times S_{(0,1)} \qquad (31)
\end{aligned}$$

, where $S_{(0,1)} = \sum_{P_2=1}^{K_2} P\left(R_{1,0,P_2} \mid H_{0,K_1,K_2}\right)$ which represents the probability that there are at least one significant stage 1 eQTL tests among the $K_2$ tests but no significant stage 1 eQTL tests among the $K_1$ tests.

The term $(29)$ demonstrates the scenario when those significant stage 1 eQTL tests present both in the $K_1$ and $K_2$ gene-by-tissue pairs. In this scenario, $\alpha^* = P\left(R^*_{2,P} \mid R_{1,P_1,P_2}, H_{0,K_1,K_2}\right)$ and therefore,

$$\begin{aligned}
&\sum_{P_1\geq 1, P_2 \geq 1} P\left(R^*_{2,P} \mid R_{1,P_1,P_2}, H_{0,K_1,K_2}\right) \times P\left(R_{1,P_1,P_2} \mid H_{0,K_1,K_2}\right) \\
&= \alpha^* \times \left\{\sum_{P_1\geq 1, P_2 \geq 1} P\left(R_{1,P_1,P_2} \mid H_{0,K_1,K_2}\right)\right\} \\
&= \alpha^* \left(1 - S_{(0,0)} - S_{(0,1)} - S_{(1,0)}\right), \qquad (32)
\end{aligned}$$

where $S_{(0,0)}$ represent the probability that under $H_{0,K_1,K_2}$, there are no significant stage 1 eQTL tests among the $K$ gene-by-tissue pairs. Note that

$S_{(0,1)} = P($ there are at least one significant stage 1 tests among the $K$ tests $\mid H_{0,K_1,K_2})$

$-P($there are at least one significant stage 1 tests among the $K_1$ tests $\mid H_{0,K_1,K_2})$

$\geq 1 - S_{(0,0)} - \alpha,$

and therefore

$$\alpha \geq 1 - S_{(0,0)} - S_{(0,1)} \,. \tag{33}$$

We define $\delta$ to be $\alpha^* - \alpha$, which represents the difference between the empirical and nominal false positive rates of the SS2 stage 2 test without applying the stage 1 test. We combine the results in (30), (31), (32) and (33) to get

$$\begin{aligned} P\left(R_{SS2}^* \mid H_{0,K_1,K_2}\right) &\leq \alpha S_{(0,1)} + \alpha^* S_{(1,0)} + \alpha^*\left(1 - S_{(0,0)} - S_{(1,0)} - S_{(0,1)}\right) \\ &= \alpha S_{(0,1)} + \alpha^*\left(1 - S_{(0,0)} - S_{(0,1)}\right) \\ &= \alpha S_{(0,1)} + (\alpha + \delta)\left(1 - S_{(0,0)} - S_{(0,1)}\right) \\ &= \alpha\left(1 - S_{(0,0)}\right) + \delta\left(1 - S_{(0,0)} - S_{(0,1)}\right) \\ &\leq \alpha\left(1 - S_{(0,0)}\right) + \delta \times \alpha \\ &\leq \alpha + \delta \times \alpha \\ &= \alpha \times (1 + \delta). \end{aligned} \tag{34}$$

Therefore under the $H_{0,K,h_{34}}$,

$$\begin{aligned} P\left(R_{SS2}^* \mid H_{0,K,h_{34}}\right) &= \textstyle\sum_{K_1 \geq 1, K_2 \geq 1} P\left(R_{SS2}^* \mid H_{0,K_1,K_2}\right) \times P\left(H_{0,K_1,K_2} \mid H_{0,K,h_{34}}\right) \\ &\leq \textstyle\sum_{K_1 \geq 1, K_2 \geq 1} \alpha(1 + \delta) \times P\left(H_{0,K_1,K_2} \mid H_{0,K,h_{34}}\right) \\ &\leq \alpha \times (1 + \delta). \end{aligned} \tag{35}$$

So far, we have shown that when the $K$ tests consist of a mixture of $H_{03}$ and $H_{04}$, the upper bound is $\alpha \times (1 + \delta)$, which could be bigger than $\alpha$. This upper bound works for any value of $K$, which would not change as the number of tests increases.

Taking into consideration all of the scenarios, the FWER of the SS2 in (21) and (22) can be calculated by

$$\begin{aligned}
P\left(R_{SS2}^{*} \mid H_{0,K}\right) &= P\left(R_{SS2}^{*} \mid H_{0,K,nsp}\right) \times P\left(H_{0,K,nsp} \mid H_{0,K}\right) \\
&+\left\{P\left(R_{SS2}^{*} \mid H_{0,K,h_4}\right) \times P\left(H_{0,K,h_4} \mid H_{0,K,sp}\right)\right. \\
&+P\left(R_{SS2}^{*} \mid H_{0,K,h_3}\right) \times P\left(H_{0,K,h_3} \mid H_{0,K,sp}\right) \\
&\left.+P\left(R_{SS2}^{*} \mid H_{0,K,h_{34}}\right) \times P\left(H_{0,K,h34} \mid H_{0,K,sp}\right)\right\} \times P\left(H_{0,K,sp} \mid H_{0,K}\right) \\
&\le \alpha \times \left\{P\left(H_{0,K,nsp} \mid H_{0,K}\right) + P\left(H_{0,K,h_4} \mid H_{0,K}\right) + P\left(H_{0,K,h_3} \mid H_{0,K}\right)\right\} \\
&+\alpha(1+\delta) \times P\left(H_{0,K,h_{34}} \mid H_{0,K}\right) \\
&\le \alpha \times (1+\delta).
\end{aligned}$$

If we want to strictly control the upper bound of the FWER at a certain level such as 0.05, $\alpha \times (1+\delta) = 0.05$, where $\delta = \alpha^{*} - \alpha$. This crude upper bound can be used to find the nominal level for performing the test in practice. By assuming a maximum value of $\alpha^{*}$ to be 1, we can solve $\alpha = 0.0253$. However, this procedure is found to be unnecessary from our empirical studies, because $\alpha^{*}$ never approaches 1 and we did not observe a single iteration with the FWER greater than the specified significance level without this procedure.

### Supplemental References


1. Zhu, Z., Zhang, F., Hu, H., Bakshi, A., Robinson, M.R., Powell, J.E., Montgomery, G.W., Goddard, M.E., Wray, N.R., Visscher, P.M., et al. (2016). Integration of summary data from GWAS and eQTL studies predicts complex trait gene targets. Nature Genetics 48, 481-487.
2. Wu, Y., Zeng, J., Zhang, F., Zhu, Z., Qi, T., Zheng, Z., Lloyd-Jones, L.R., Marioni, R.E., Martin, N.G., Montgomery, G.W., et al. (2018). Integrative analysis of omics summary data reveals putative mechanisms underlying complex traits. Nature Communications 9, 918.
3. Giambartolomei, C., Vukcevic, D., Schadt, E.E., Franke, L., Hingorani, A.D., Wallace, C., and Plagnol, V. (2014). Bayesian Test for Colocalisation between Pairs of Genetic Association Studies Using Summary Statistics. PLOS Genetics 10, e1004383.
4. Dobbyn, A., Huckins, L.M., Boocock, J., Sloofman, L.G., Glicksberg, B.S., Giambartolomei, C., Hoffman, G.E., Perumal, T.M., Girdhar, K., Jiang, Y., et al. (2018). Landscape of Conditional eQTL in Dorsolateral Prefrontal Cortex and Co-localization with Schizophrenia GWAS. Am J Hum Genet 102, 1169-1184.

5. Lonsdale, J., Thomas, J., Salvatore, M., Phillips, R., Lo, E., Shad, S., Hasz, R., Walters, G., Garcia, F., Young, N., et al. (2013). The Genotype-Tissue Expression (GTEx) project. Nature Genetics 45, 580-585.

6. Ongen, H., Buil, A., Brown, A.A., Dermitzakis, E.T., and Delaneau, O. (2016). Fast and efficient QTL mapper for thousands of molecular phenotypes. Bioinformatics 32, 1479-1485.

7. Gong, J., Wang, F., Xiao, B., Panjwani, N., Lin, F., Keenan, K., Avolio, J., Esmaeili, M., Zhang, L., He, G., et al. (2019). Genetic association and transcriptome integration identify contributing genes and tissues at cystic fibrosis modifier loci. PLOS Genetics 15, e1008007.

8. Corvol, H., Blackman, S.M., Boëlle, P.-Y., Gallins, P.J., Pace, R.G., Stonebraker, J.R., Accurso, F.J., Clement, A., Collaco, J.M., Dang, H., et al. (2015). Genome-wide association meta-analysis identifies five modifier loci of lung disease severity in cystic fibrosis. Nature Communications 6, 8382.

9. Hormozdiari, F., Kostem, E., Kang, E.Y., Pasaniuc, B., and Eskin, E. (2014). Identifying causal variants at loci with multiple signals of association. Genetics 198, 497-508.

10. Bulik-Sullivan, B.K., Loh, P.-R., Finucane, H.K., Ripke, S., Yang, J., Patterson, N., Daly, M.J., Price, A.L., and Neale, B.M. (2015). LD Score regression

distinguishes confounding from polygenicity in genome-wide association studies. Nature genetics 47, 291-295.

11. Xu, Z., Duan, Q., Yan, S., Chen, W., Li, M., Lange, E., and Li, Y. (2015). DISSCO: direct imputation of summary statistics allowing covariates. Bioinformatics 31, 2434-2442.
12. Hormozdiari, F., van de Bunt, M., Segrè, A.V., Li, X., Joo, J.W.J., Bilow, M., Sul, J.H., Sankararaman, S., Pasaniuc, B., and Eskin, E. (2016). Colocalization of GWAS and eQTL Signals Detects Target Genes. Am J Hum Genet 99, 1245-1260.
13. Baltagi, B.H. (2008). Econometrics.(Berlin, Heidelberg: Springer Berlin Heidelberg).
14. Park, J.Y., Wu, C., Basu, S., McGue, M., and Pan, W. (2018). Adaptive SNP-set association testing in generalized linear mixed models with application to family studies. Behavior genetics 48, 55-66.
15. Park, J.Y., Wu, C., and Pan, W. (2018). An adaptive gene-level association test for pedigree data. BMC genetics 19, 39-43.
16. Laird, N.M., and Ware, J.H. (1982). Random-effects models for longitudinal data. Biometrics, 963-974.

17. Lindstrom, M.J., and Bates, D.M. (1988). Newton—Raphson and EM algorithms for linear mixed-effects models for repeated-measures data. Journal of the American Statistical Association 83, 1014-1022.

18. Pinheiro, J., Bates, D., DebRoy, S., Sarkar, D., Heisterkamp, S., Van Willigen, B., and Maintainer, R. (2017). Package 'nlme'. Linear and nonlinear mixed effects models, version 3.

19. Chen, H., Conomos, M.P., and Chen, M.H. (2019). Package 'GMMAT'.
